# Diabetic phenotype in mouse and humans with β-amyloid pathology reduces the number of microglia around β-amyloid plaques

**DOI:** 10.1101/2020.04.07.027292

**Authors:** Teemu Natunen, Henna Martiskainen, Mikael Marttinen, Sami Gabbouj, Hennariikka Koivisto, Susanna Kemppainen, Satu Kaipainen, Mari Takalo, Helena Svobodová, Luukas Leppänen, Benjam Kemiläinen, Simo Ryhänen, Teemu Kuulasmaa, Eija Rahunen, Sisko Juutinen, Petra Mäkinen, Pasi Miettinen, Tuomas Rauramaa, Jussi Pihlajamäki, Annakaisa Haapasalo, Ville Leinonen, Heikki Tanila, Mikko Hiltunen

## Abstract

Type 2 diabetes (T2D) increases the risk of Alzheimer’s disease (AD). Even though these two diseases share common molecular pathways, the mechanisms remain elusive. To shed light into these mechanisms, mice with different AD- and/or tauopathy-linked genetic backgrounds were utilized; APPswe/PS1dE9 (A+Tw), Tau P301L (AwT+), and APPswe/PS1dE9/Tau P301L (A+T+). Feeding these mice with typical Western diet (TWD) led to obesity and diabetic phenotype as compared to respective mice with a standard diet. TWD also exacerbated memory and learning impairment in A+Tw and AwT+, but not in A+T+ mice. Furthermore, RNA sequencing of mouse hippocampal samples revealed altered responses to AD-related pathologies in A+Tw and A+T+ mice upon TWD, pointing specifically towards aberrant microglial functionality and PI3K-Akt signaling. Accordingly, fewer microglia alongside an increased number of dystrophic neurites around β-amyloid plaques, and impaired PI3K-Akt signaling, were discovered in the hippocampus of TWD mice. Mechanistic elucidation revealed that disruption of the PI3K-Akt signaling pathway by pharmacological or genetic approaches significantly decreased the phagocytic uptake and proinflammatory response as well as increased the activity of Syk-kinase upon ligand-induced activation of Trem2/Dap12 signaling in mouse microglia. Finally, characterization of microglial pathology in cortical biopsies of idiopathic normal pressure hydrocephalus (iNPH) patients harboring β-amyloid plaques revealed a significant decrease in the number of microglia per β-amyloid plaque in obese iNPH patients with T2D as compared to both normal weight and obese iNPH patients without T2D. Collectively, these results suggest that the peripheral diabetic phenotype in mice and humans associates with reduced microglial response to β-amyloid pathology.

## Introduction

Alzheimer’s disease (AD) is the most common neurodegenerative disease. Typical AD pathology includes accumulation of neuritic plaques composed of β-amyloid peptide (Aβ) aggregates, and dystrophic neurites positive for hyperphosphorylated Tau in the form of neurofibrillary tangles (NFTs) [6, 64]. Furthermore, activation of microglia, loss of synapses, and neuronal death are typically observed in AD [28]. Currently, available AD medication offers only a minor symptomatic relief, without any effects on the underlying pathogenic processes. Although AD has a strong genetic background, environmental and lifestyle factors, such as type 2 diabetes (T2D), play important roles in conferring the risk for AD [31]. T2D is characterized by insulin resistance in peripheral tissues, leading to increased serum insulin and glucose levels. Interestingly, AD and T2D share common features, such as impaired insulin signaling, increased levels of pro-inflammatory cytokines, metabolic changes, increased oxidative stress, and mitochondrial dysfunction [16]. However, the exact common underlying mechanisms between these two diseases remain elusive. Continuously increasing prevalence of both AD and T2D world-wide warrants for a better mechanistic understanding of the contribution of T2D on AD risk.

Lifestyle factors, such as lack of exercise and typical Western diet (TWD) containing high fat, high sugar, high cholesterol, and low fiber play a crucial role in the development of T2D. Likewise, epidemiological studies have revealed that the consumption of saturated fats, T2D, and obesity in midlife increase the risk of AD later in life [14, 21, 31, 45, 50, 103]. In mice, high-fat diet (HFD) and TWD lead to obesity, glucose intolerance and finally to full-blown T2D [101]. In most of the studies, HFD and TWD have led to impaired memory in both wild-type (WT) and AD transgenic mice [24, 26, 47, 81]. Also, impaired insulin-Akt signaling in the brain of HFD/TWD animals has been often reported [34, 48, 73]. However, effects of HFD/TWD on β-amyloid and Tau pathology have been controversial [22, 24, 26, 36, 99].

In the brain, insulin plays a role as a neurotrophic factor and has also been linked to neuroinflammation, while neuronal glucose intake is mainly independent of the insulin-regulated GLUT-4 transporter [12, 108]. When insulin binds to insulin receptor (IR), it can activate two distinct branches of insulin signaling pathway: the Ras-mitogen-activated protein kinase (MAPK) and phosphoinositide 3-kinase (PI3K)-Akt-kinase pathways [41]. Akt-kinase controls the activity of GSK3β. Reduced levels of GSK3β phosphorylation, specifically at the inhibitory S9 residue, in turn, result in augmented Tau phosphorylation [30]. Importantly, Akt-signaling also controls autophagic activity through mammalian target of Rapamycin complex 1 (mTORC1). Impaired autophagy is a common feature in AD and other neurodegenerative diseases [89]. Activation of PI3K-Akt-mTORC1 pathway by insulin or other stimuli inhibits autophagy, while nutrient starvation inactivates this pathway, and thus increases autophagic activity [25].

Inflammation is one of the key elements shared in both AD and T2D. Increased levels of proinflammatory cytokines, including tumor necrosis factor α (TNFα), are observed in adipose tissue of both obese and diabetic subjects as well as in the brain of AD patients [88, 93]. Recent genetic and functional studies have highlighted the relevance of the brain’s innate immune cells in contributing to the onset and progression of AD [69, 87]. Specifically, studies have exemplified the emergence of reactive microglia subtypes in response to disease and damage, for which triggering receptor expressed on myeloid cells 2 (Trem2) has been identified as a central receptor [40, 49]. Transition of microglia from homeostatic to a disease-associated microglia (DAM) phenotype is a two-step process, in which Trem2-PI3K-Akt pathway plays a central role [40]. Furthermore, the deficits of Trem2 impair Akt-mTOR signaling in microglia, affecting autophagy and decreasing the ability of microglia to form a protective barrier around β-amyloid-plaques, which eventually leads to increased formation of dystrophic neurites [95].

Here, we have studied the interaction between dietary (TWD) and genetic factors (APPswe/PS1dE9 and Tau P301L mutations) and their effects on memory, brain pathology, and global gene expression in female mice with WT (AwTw) and different AD- and tauopathy-linked genetic backgrounds: Tau-P301L (AwT+), APPswe/PS1dE9 (A+Tw), and triple transgenic APPswe/PS1dE9/Tau-P301L (A+T+). To our knowledge, this is the first time when all these mouse lines have been utilized in parallel in the same study to dissect the alterations in cellular pathways in the brain associated with TWD and different AD-associated pathologies. Our results suggest that TWD exacerbates age-related memory impairment in mice with a genetic predisposition to develop AD-like brain pathology as well as alters microglial gene expression and functionality. Supporting these findings, T2D also decreased the number of β-amyloid plaque-associated microglia also in frontal cortical biopsies of living idiopathic normal pressure hydrocephalus (iNPH) patients.

## Materials and Methods

### Animals

In this study, we used three transgenic mouse lines with C57Bl/6J background as models for AD and WT littermates with the same C57Bl/6J background as controls. The first mouse line coexpressed in a single transgene a chimeric mouse/human APP695 harboring the Swedish K670M/N671L mutations (Mo/Hu APPswe) and human PS1 with the exon-9 deletion (PS1dE9) under mouse PrP promoter [32]. The second mouse line carried human P301L tau mutation under CaMKII promoter [43]. The third line was created by crossing heterozygous line 1 and 2 mice resulting in APPswe/PS1dE9 x P301Ltau mice. The study comprised of 46 female mice, 11 WT (AwTw), 13 APPswe/PS1dE9 transgenic (A+Tw), 11 tau P301L transgenic (AwT+) and 11 transgenic for both APPswe/PS1dE9 and tau P301L (A+T+). The dietary intervention started at the age of 7 months, so that half of the mice in each genotype group received a chow mimicking TWD (Adjusted Calories diet, TD 88137, Harlan Tecklad, Madison, WI, USA, with 21% w/w fat, 0.15 % cholesterol and 35% sucrose) while the other half continued with standard rodent diet (STD) (5% w/w fat and 0% cholesterol) until sacrificed at the age of 13 months. The mice were weighed monthly. The mice were kept in a controlled environment (constant temperature, 22 ± 1 °C, humidity 50 – 60 %, lights on 07:00-19:00), with food and water available ad libitum. All animal procedures were carried out in accordance with the guidelines of the European Community Council Directives 86/609/EEC and approved by the Animal Experiment Board of Finland.

### Glucose tolerance test (GTT)

GTT for mice was done at the age of 11 months. After 3 h of fasting in the morning, an i.p. injection of 1 mg/g D-glucose in a 20% solution (prepared in normal saline) was given. Blood samples for the determination of glucose levels (50 – 75 µl) were collected at time points 0 (before glucose injection) and 30 min after the injection from the saphenous vein. The glucose values were determined immediately using a glucometer (One Touch, LifeScan Inc., Milpitas, CA, USA).

### Spontaneous activity

Spontaneous explorative activity was assessed by using an automated activity monitor (TruScan, Coulbourn Instruments, Whitehall,PA, USA) based on infrared photobeam detection. The system consisted of an observation cage with white plastic walls (26 cm x 26 cm x 39 cm) and two frames of photo detectors enabling separate monitoring of horizontal (XY-move time) and vertical activity (rearing). The test cage was cleaned with 70% ethanol before each mouse to avoid odor traces. The following parameters were measured during a 10-min session: ambulatory distance (gross horizontal locomotion) and rearing time.

### Morris swim navigation test

Spatial learning and memory were assessed in the Morris swim task. The test was conducted in a white circular wading pool (diameter 120 cm) with a transparent submerged platform (diameter 14 x 14 cm) 1.0 cm below the surface serving for escape from the water. The pool was open to landmarks in the room (white screen blocking the view to the computer and the experimenter, green water hose, door, 1-m high black pattern on the wall). Temperature of the water was kept at 20 ± 0.5°C. The acquisition phase was preceded by two practice days with a guiding alley to the platform (day −4 and day −3, not shown). During the acquisition phase (days 1-5), the location of the hidden platform was kept constant (SE quadrant) and the starting position varied between four different locations at the pool edge, with all mice starting from the same position in a given trial. Each mouse was placed in the water with its nose pointing towards the pool wall. If the mouse failed to find the escape platform within 60 s, it was placed on the platform for 10 s by the experimenter (the same time was allowed for mice that found the platform). The acquisition phase consisted of five daily trials with a 10 min inter-trial-interval. On day 5, the search bias was tested in a 60-s probe trial (the 5th trial) without the platform. The experimenter was blind to the genotype and treatment of the mice. The mouse was video-tracked, and the video analysis program calculated the escape latency, swimming speed, path length and time in the pool periphery (10 cm from the wall) and in the platform zone (diameter 30 cm).

### Passive avoidance test

This task was used as a control for long-term memory, since in contrast to Morris swim task, increased activity level leads to impaired performance in this task. The mouse was placed in the well-lit side of a two-compartment box and was freely allowed to enter the dark, closed compartment through a hole in the dividing wall. As soon as the mouse entered the dark side, the slide door separating the compartment was closed and a mild foot-shock delivered (2 x 2 s at 0.30 mA). The mouse was then taken to its home cage. The memory for the aversive experience was assessed 48 h later by taking the time for the mouse to enter the dark compartment with a cut-off time of 180s.

### Tissue sampling and preparation

At the end of the study, all mice were deeply anesthetized with intraperitoneal pentobarbiturate-chloralhydrate cocktail (60 mg/kg each) and transcardially perfused with ice-cold saline for 3 min to rinse blood from the brain. The brain was removed and placed on ice. For half of the mice, one brain hemisphere was immersion fixed in 4% paraformaldehyde, followed by 30 % sucrose overnight and stored in antifreeze at – 20°C. The other hemisphere was dissected on ice into following blocks: frontal, parieto-occipital and temporo-occipital cortices, hippocampus, cerebellum and olfactory bulb. In addition, two lobes of liver, pancreas, gastrocnemius and tibialis anterior muscles and samples of inguinal and subcutaneous fat were dissected and snap frozen in liquid nitrogen. The samples were stored at −80°C. The PFA 4% fixed hemispheres were cut using a freezing slide microtome into 35 µm coronal sections. The hippocampal tissue samples were collected into microcentrifuge tubes and weighed. Samples were homogenized in 250 µl of phosphate-buffered saline (DPBS, Lonza, Walkersville, MD, USA) using a stirrer. Fractions of homogenates were taken to RNA isolation (50 µl of homogenate and 500 µl Tri Reagent (Sigma-Aldrich, St. Louis, MO, USA) and Western blot analysis (100 µl of homogenate was supplemented with EDTA-free protease inhibitor cocktail (ThermoScientific, Waltham, MA, USA) and HALT™ phosphatase inhibitor cocktail (ThermoScientific, Waltham, MA, USA) 1:100. The remaining 100 µl of homogenate was left unprocessed and stored at −80°C.

### Histochemistry

Snap frozen liver blocks (originally designed for biochemistry) were cut with a cryostat into 5 µm sections and stained with hematoxylin-eosin to reveal lipid vacuoles indicating a fatty liver change. Despite the freezing artefact, the vacuoles were clearly visible in most cases. Three sections of each mouse were analyzed by two researchers blinded to the genotype and treatment of the mice and scored as follows: 0 = no change, 1 = possible change, 2 = clear fatty-liver change. The brain hemispheres fixed in 4% paraformaldehyde were cut using a freezing slide microtome into 35 µm coronal sections. Three brain sections, 105 µm apart were selected between bregma −3.1 mm and - 3.5 mm coronal planes according to the Paxinos and Franklin atlas [71]. IHC staining was done with fluorescent secondary antibodies for free-floating sections. First, the sections were pre-treated in 0.05M citrate solution for 30 min (pH 6.0) at 80⏑. Endogenous peroxidase activity on sections was inhibited by incubation with 0.3% or 2% hydrogen peroxide in methanol. Non-specific antibody binding was blocked with 3% bovine serum albumin or 10 % normal goat serum. To visualize β-amyloid burden and dystrophic neurites, the sections were stained with rabbit polyclonal anti-Pan-Aβ (1:2000, Aβ 1-40, Invitrogen, Carlbad, CA, USA) and mouse monoclonal antihuman phospho-Tau (S202, T205) AT8 (1:1000, Thermo Fisher Scientific, MN1020, UK) for dystrophic neurites. To see details of microglia around β-amyloid plaques, sections were stained with two sets of antibodies. β-amyloid plaques were stained with X-34 (SML1954, Sigma-Aldrich, St. Louis, MO, USA), microglia with rabbit polyclonal anti-Iba-1 (1:5000, #019-19741, FUJIFILM Wako Chemicals Europe GmbH, Neuss, Germany) and lysosomes with rat monoclonal anti-Cd68 (clone FA-11, #MCA1957, Bio-Rad, Hercules, CA, USA). In the second set, p85 was stained with mouse monoclonal anti-p85 (1:1000, Antibodies-online GmbH, Aachen, Germany) along with Iba-1 staining. Primary antibodies were incubated overnight at +4⏑. For X-34 staining, the sections were incubated in X-34 solution for 1 h and briefly rinsed with 60%⏑[PBS / 40%⏑[EtOH solution prior to primary antibody incubation. The sections were incubated with appropriate fluorescent secondary antibodies: goat anti-mouse 488 (Invitrogen Alexa Fluor 488, A11029, Molecular Probes, Invitrogen, Eugene, OR, USA) for p85 and AT8, goat anti-rabbit 594 (Invitrogen Alexa Fluor 594, A11037, Molecular Probes, Invitrogen, Eugene, OR, USA) for Iba-1 and Pan-Aβ, and goat ant-rat 488 (Invitrogen Alexa Fluor 488, A1106, Molecular Probes, Invitrogen, Eugene, OR, USA) for Cd68. The sections were mounted with Vectashield Hard Set with or without DAPI (Vector Laboratories, Burlingame, CA, USA) on gelatin-coated slides. Control sections without the primary antibodies were processed simultaneously, and no unspecific staining was observed.

### Fluorescent microscopy and image analysis

The analysis of β-amyloid plaques and surrounding dystrophic neurites was done in the dentate gyrus (DG) and lateral entorhinal cortex (LEC) (Fig. 1I). Images were taken with a fluorescent microscope (Zeiss Axio Imager M2 microscope, Germany, with AxioCam ERc5s -camera attached). Both brain areas were imaged with 5x or 20x objectives. Brain areas were imaged with Z-stack imaging method, 12 layers with a total distance of 15 µm. Dystrophic neurite counting was done with direct microscopy. The ZEN 2012 software Blue edition (Carl Zeiss Microimaging GmbH) was used for image processing. The β-amyloid plaque burden was determined with Adobe Photoshop CS6 extended (version 13.0×32). To estimate the total β-amyloid burden in the studied brain regions, we measured the total area of β-amyloid plaques with the most representative threshold, same for all animals, within the defined ROI (LEC and HC) in three brain sections and divided the area by the ROI area. Thus, the β-amyloid burden was expressed as an average percentage from three brain sections. All plaques over 9.9 µm diameter in both brain areas, LEC and HC, were counted as well as dystrophic neurites around each plaque. Counting was done by an investigator blinded to the genetic and dietary background of mice.

**Fig. 1.**
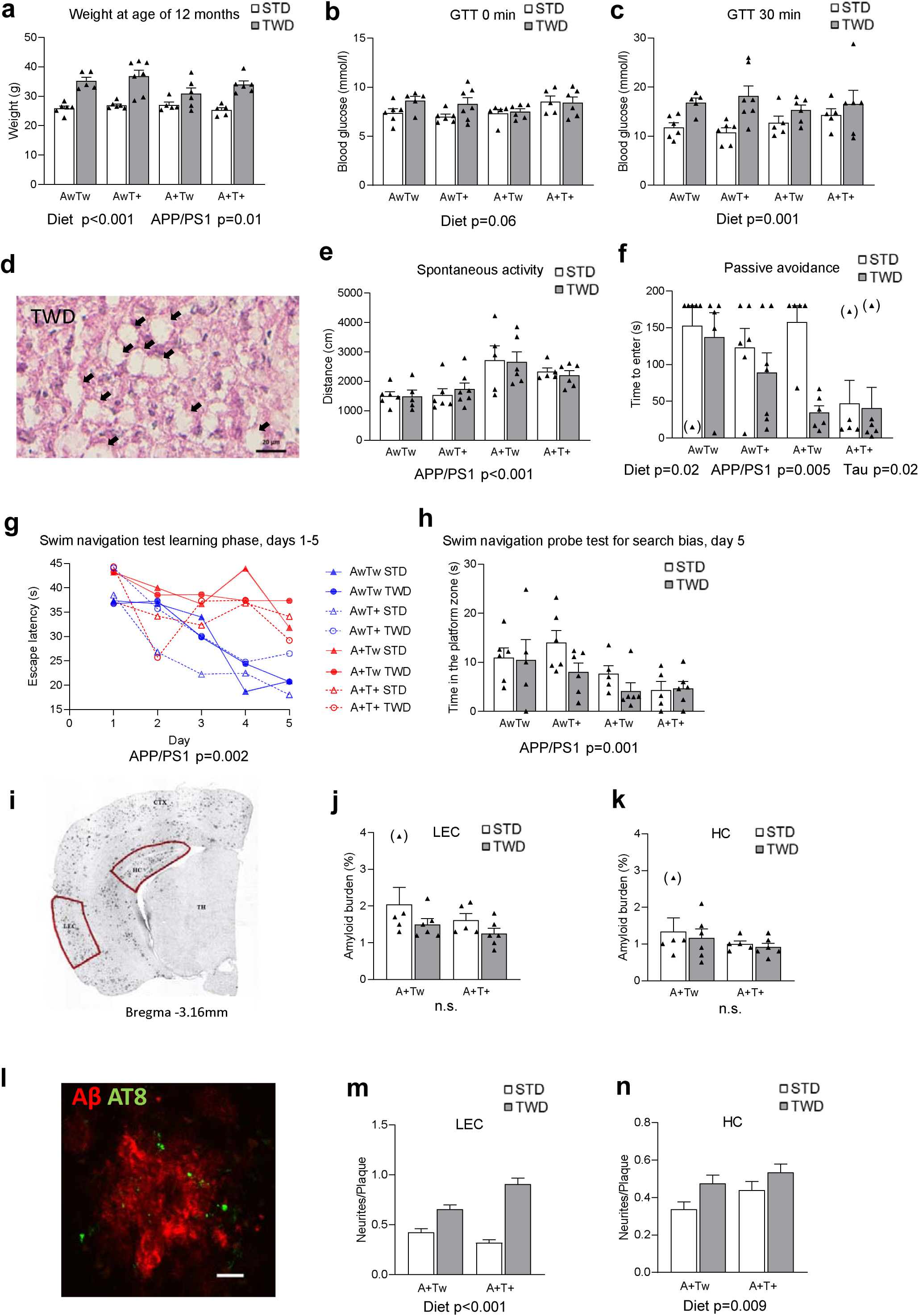
TWD leads to T2D, memory impairment and increased number of dystrophic neurites in mice with AD-linked genetic backgrounds. **a** Weight of mice at age of 12 months (Two-way ANOVA). **b** Glucose tolerance test (GTT) results after 3h fasting (Two-way ANOVA). **c** Serum glucose levels in GTT after 30 min of D-glucose injection (p=0.001, Two-way ANOVA). **d** Representative liver sample image of TWD. Mice with TWD had significantly higher fatty liver score as compared to mice with STD (p<0.001, Mann-Whitney). Black arrows indicate lipid vacuoles. Scale bar 20µm. **e** Assessment of spontaneous activity (p<0.001, Two-way ANOVA). **f** Passive avoidance test revealed impaired performance of mice with TWD as compared to mice with STD (diet effect, p=0.02, Two-way ANOVA). Also AD and tauopathy-associated transgenes impaired the performance (APPswe/PS1dE9; p<0.001 and Tau; p<0.001, Two-way ANOVA). Three outliers were left out from statistical analysis (in brackets) **g** In five-day learning phase of Morris swim navigation test, mice with APPswe/PS1dE9 transgene (red lines) did not learn to find the platform as fast as mice without APPswe/PS1dE9 transgene (blue lines, p=0.002, Two-way ANOVA), while TWD and Tau transgene did not affect learning. **h** On day five, mice with APPswe/PS1dE9 transgene spent significantly less time in platform zone (p=0.001, Two-way ANOVA). Also, mice with TWD spent less time in platform zone as compared to mice with STD, but the difference did not reach statistical significance (p<0.09, Two-way ANOVA). **i** A representative image of APPswe/PS1dE9 mouse coronal brain section used for assessing β-amyloid plaque load from lateral entorhinal cortex (LEC) and from dentate gyrus (DG) of hippocampus (HC). **j** Amyloid burden quantification in LEC, and **k** in HC. **l** A representative fluorescence microscope image of APPswe/PS1dE9 mouse coronal brain section showing AT8-positive dystrophic neurites around β-amyloid plaques. **m** Quantification of neurites/plaque in LEC showing significant increase in TWD mice as compared to STD mice (p<0.001). **n** Also, in HC number of neurites per plaque was increased in TWD mice as compared to STD mice (p=0.009). All results are shown as mean + SEM, behavioral tests: n=5-7 mice/group, Two-way ANOVA, immunohistochemistry: n=5-6 mice/group, 3 brain slices/mouse. Dystrophic neurites around β-amyloid plaques (diameter >10μM) were counted, Kruskal-Wallis H-test.

### Confocal microscopy and image analysis

Details of microglia around β-amyloid plaques were analyzed from fluorescent images of the molecular layer of dentate gyrus obtained with a Zeiss Axio Observer inverted microscope (63 × NA 1.4 oil objective) equipped with a Zeiss LSM 700 or Zeiss LSM 800 confocal module (Carl Zeiss Microimaging GmbH, Jena, Germany). For all analyses, at least three z-stack images (9 µm with 0.9 µm optical sections) were collected per brain section. Laser and detector settings were maintained constant for each immunostaining. All image processing and analyses were performed with Fiji software [84] using automated scripts. Maximum intensity projections were generated for each channel, and Gaussian blurring and rolling ball algorithm were applied to remove noise and subtract background, respectively. The stainings were segmented using Otsu threshold for X-34, and Moments threshold for Iba1, Cd68, and p85. For the analysis of Cd68 and Iba1, the outlines of a β-amyloid plaque were first determined automatically from the X-34 staining, then Cd68 and Iba1 staining area and intensity were analyzed within 30 µm from the plaque outline. When analyzing p85, outlines of β-amyloid plaque was assessed manually due to a lack of X-34 staining, and then p85 staining area and intensity were analyzed within 30 µm from the plaque outline. Imaging and analysis of the immunostainings was done by investigators blinded to the genetic or dietary background of mice.

### iNPH cohort for cortical brain biopsies

Patients characterized in Table 1, were diagnosed with probable iNPH [78] and selected for CSF shunt surgery according to the previously described protocol [37] in Department of Neurosurgery Kuopio University Hospital from 2012 until 2017. Diagnostic right frontal cortical brain biopsy was taken during shunt surgery and occurrence of β-amyloid and/or phospho-Tau was evaluated by neuropathologist as described previously [58]. According to the neuropathological examination, all the individuals included in this study were β-amyloid positive but phospho-Tau (AT8) negative. All patients gave their written informed consent, prior to the surgery, for the study accepted by Ethics Committee, Hospital District of Northern Savo.

**Table 1.** Demographics, clinical status, and pathology of iNPH cases

| Group | Gender | Age<br>(years) | Weight<br>(kg) | Height<br>(cm) | BMI<sup>(1)</sup> | HOMA-IR<sup>(2)</sup> | T2D<sup>(3)</sup> | Hyper-<br>tension | Aβ<br>pathology | APOE<br>genotype |
| --- | --- | --- | --- | --- | --- | --- | --- | --- | --- | --- |
| **BMI<25** | Female | 67 | 70 | 170 | 24.2 | 0.6 | no | no | positive | 34 |
|  | Male | 79 | 65 | 168 | 23 | 2.7 | no | no | positive | 34 |
|  | Male | 75 | 56 | 159 | 22.2 | 0.7 | no | yes | positive | 44 |
|  | Female | 71 | 60 | 158 | 24 | 1.9 | no | no | positive | 33 |
|  | avg. | 73 | 63 | 164 | 23.4 | 1.5 |  |  |  |  |
| **BMI>30** | Female | 71 | 87 | 160 | 34 | 4.9 | no | no | positive | 34 |
|  | Male | 81 | 102 | 171 | 34.9 | 7 | no | no | positive | 33 |
|  | Male | 69 | 80 | 160 | 31.3 | 3.4 | no | yes | positive | 33 |
|  | avg. | 73.5 | 89.8 | 166 | 32.6 | 5.1 |  |  |  |  |
| **BMI>30 + T2D** | Male | 72 | 98 | 170 | 33.9 | 14.1 | yes | yes | positive | 33 |
|  | Male | 73 | 90 | 173 | 30.1 | 4.7 | yes | yes | positive | 34 |
|  | Male | 73 | 92 | 162 | 35.1 | 4.5 | yes | yes | positive | 33 |
|  | Female | 75 | 92 | 155 | 38.3 | 16.8 | yes | yes | positive | 34 |
|  | Male | 67 | 82 | 165 | 30.1 | 5.9 | yes | yes | positive | 34 |
|  | avg. | 72 | 90.8 | 165 | 33.5 | 9.2 |  |  |  |  |

Age, Weight, Height, BMI and clinical status when biopsy was taken
<sup>1)</sup> Body Mass Index, weight (kg) / [height (m)]<sup>2</sup>
<sup>2)</sup> Homeostatic Model Assessment for Insulin Resistance = Blood Insulin ( $\mu$ U/l) x Glucose (mg/dl)
<sup>3)</sup> Type 2 Diabetes

### Immunohistochemistry of human biopsy brain tissue

The biopsy samples were fixed in buffered formalin overnight and subsequently embedded in paraffin. 5 µm deparaffinized sections were used for IHC staining. Antigen retrieval was performed by boiling the sections in 10 mM Tris – 1 mM EDTA buffer, pH 9.0, in pressure cooker for 10 minutes, followed by cooling to room temperature and incubation in 80 % formic acid for 20 minutes. Endogenous peroxidase activity was quenched by incubation in 1 % H_2_O_2_ and unspecific antibody binding was blocked using 3 % bovine serum albumin (BSA). Sections were incubated with the first primary antibody, rabbit polyclonal anti-Iba-1 (#019-19741, 1:1500, FUJIFILM Wako Chemicals Europe GmbH, Neuss, Germany) overnight at 4°C, followed by visualization with biotinylated anti-rabbit secondary antibody (1:200, Vector Laboratories, Burlingame, CA, USA) and Vectastain Elite ABC Kit Peroxidase (Vector Laboratories, Burlingame, CA, USA) using 3,3’-diaminobenzidine (DAB, Sigma-Aldrich, St. Louis, MO, USA) as chromogen with 1min 30sec incubation time. Staining of the second antigen started with blocking with 3 % BSA and incubation with the second primary antibody, mouse monoclonal anti-Aβ (anti-β-amyloid 17-24, clone 4G8, #800712, 1:500, Biolegend, San Diego, CA, USA) overnight at 4°C. Staining was visualized with biotinylated anti-mouse secondary antibody (1:200, Vector Laboratories, Burlingame, CA, USA) and Vectastain ABC-AP Kit (Vector Laboratories, Burlingame, CA, USA), and 30-minute incubation with Permanent AP Red Kit substrate/chromogen (Zytomed Systems GmbH, Berlin, Germany). Sections were counterstained with Mayer’s haematoxylin and mounted with aqueous mounting medium (Aquatex, Merck, Darmstadt, Germany). Control sections without the primary antibodies were processed simultaneously, and no unspecific staining was observed.

### Imaging and analysis of human brain biopsy tissue

The human biopsy brain tissue sections were imaged with Hamamatsu NanoZoomer-XR Digital slide scanner with 20x (NA 0.75) objective (Hamamatsu Photonics K.K.). Due to the small size of the biopsy specimen, all plaques with microglia within or around the plaque in the grey matter area of the specimen were included in the analysis. Size of the β-amyloid plaques was obtained by manually outlining the plaques in NDP.view2 software (Hamamatsu Photonics K.K.). Plaque-associated microglia were manually counted, and the number of microglia within the plaque area and around the plaque area were recorded separately. Only microglia with clearly visible soma were included in the count. Counting and analysis was performed by an investigator blinded to sample identity.

### Cell cultures and treatments

Immortalized mouse microglial BV2 cells were grown in RPMI-1640 medium (Sigma-Aldrich, St. Louis, MO, USA) supplemented with 10% fetal bovine serum (FBS, Gibco, Waltham, MA, USA), 2mM L-glutamine (Lonza, Basel, Switzerland) and 1% penicillin-streptomycin (P/S: 100 U/ml penicillin and 100 U/ml streptomycin, Lonza, Basel, Switzerland) in 15 cm cell culture plates (Nunc, Roskilde, Denmark) as previously described [23]. To study the effects of PI3K inhibition on Syk phosphorylation, BV2 cells were scraped from confluent plate on RPMI-1640 medium without FBS to induce serum starvation. Cells were diluted to concentration 1 × 10^6^ cells / ml, and cell suspension was divided into 1,5 ml Eppendorf tubes. After 1 h, PI3K inhibitor treatment with 1µM LY294002 (Life Technologies, Frederick, MD, USA) was started. After 1h 40 min, Src-kinase inhibitor (2 µM SU 6656, S9692 Sigma-Aldrich, St. Louis, MO, USA) was added. At 2 h time point, cells were treated with M-CSF (100ng/ml) Recombinant Human M-CSF (Macrophage-Colony Stimulating Factor, BioLegend cat. 574806, San Diego, CA, USA) for 5 min. Tubes were placed on ice and subsequently centrifuged at 10 000 x g for 1 min at +4°C. Supernatant was removed and cells were lysed with tissue protein extraction buffer (T-PER, ThermoScientific, Rockford, IL, USA) including EDTA-free protease inhibitor cocktail (ThermoScientific, Waltham, MA, USA) and HALT™ phosphatase inhibitor cocktail (ThermoScientific, Waltham, MA, USA) 1:100, and incubating on ice for 30min. The lysates were centrifuged at 16 000 x g for 10min at +4°C, the protein lysates were collected into the new tubes and stored at −20°C. Effects of PI3K inhibition on Syk phosphorylation in mouse WT primary microglia was studied by plating 100 000 cells in 96-well plate in Dulbecco’s modified Eagle’s medium (DMEM, Lonza, Verviers, Belgium) supplemented with 2mM L-glutamine (Lonza, Verviers, Belgium) and 1% P/S (Lonza, Verviers, Belgium), but without FBS in order to induce serum starvation. After 1 h, PI3K inhibitor treatment with 1µM LY294002 (Life Technologies, Frederick, MD, USA) was started. After 2 h, cells were treated with Trem2 antibody (R&D Systems, Minneapolis, MN, USA) with final concentration of 10 µg/ml. After 5 min antibody treatment, plate was placed on ice, cells were lysed and supernatants were analyzed with phosphoSyk Alpha-LISA kit (PerkinElmer, Waltham, MA, USA) according to the kit instructions. Mouse primary microglia cultures from Akt2 knock-out (KO) (Mouse B6.129P2(Cg)-Akt2tm1.1Mbb/J, The Jackson Laboratory, Ben Harbor, Maine, USA) and C57BL/6 WT mice were prepared as previously [68]. Briefly, brains of neonatal (P0–P2) mice were dissected and meninges were removed. Brain tissue was dissociated using mechanical shearing and trypsin. Cells of two brains were plated on poly-L-lysine (Sigma-Aldrich, St. Louis, MO, USA) coated T75 culture flasks and cultured in Dulbecco’s modified Eagle’s medium (DMEM, Lonza, Verviers, Belgium) supplemented with 10% FBS (Gibco, Waltham, MA, USA), 2mM L-glutamine (Lonza, Verviers, Belgium) and 1% P/S (Lonza, Verviers, Belgium). On the next day, the cells were washed three times with PBS (DPBS, Lonza, Walkersville, MD, USA) to remove cellular debris and cultured with fresh DMEM with supplements. After 7 days, mature microglia were shaken off from the astrocytic feeding layer. Microglial cells were counted, and appropriate number of cells was plated in 96 well plates and used for assessing phagocytic activity and responses to inflammatory activation.

### Phagocytosis assay

For pHrodo phagocytosis assays, BV2 cells were plated on 96-well plates (Nunc, Roskilde, Denmark) at 4,000 cells/well 16h before starting the assay. Primary microglia were cultured in 96-well plates at 20,000 cells/well in volume of 100µl in DMEM culture medium with supplements described above. Cells were treated with PI3K inhibitor LY294002 (Life Technologies, Frederick, MD, USA) with different concentrations and with 5µM Cytochalasin D (Sigma-Aldrich/Merck, St. Louis, MO, USA), inhibitor of actin polymerization, which served as negative control by blocking the phagocytosis. After 1h pretreatment with the inhibitors, cells were treated with pHrodo Zymosan bioparticles (Essen BioScience, Ann Arbor, MI, USA) with a final amount of 5µg / well. Three to four wells were used for each treatment condition. Time-lapse videomicroscopy sequences of living cells were obtained using the IncuCyte S3 system (Essen BioScience, Ann Arbor, MI, USA) with 20x (BV2) or 10x (primary microglia) objective lense, and images from 4 fields per culture well were obtained every 15 minutes (BV2) or every 30 minutes (primary microglia) for up to 3 hours. pHrodo red fluorescent signal was used as an indicator of uptake by the lysosomal cell compartments. After 3h, nuclei of cells were stained with 1 µM Vybrant DyeCycle Green (Molecular probes/ThermoFisher Scientific, Eugene, OR, USA) and imaged with IncuCyte in order to count the cells. Area (µm^2^) of red fluorescence per image was calculated for each time point, normalized to total cell count, and plotted as a time course.

### Western blot analysis

The inhibitor-supplemented total protein fractions were further diluted by taking 50 µl of homogenate and adding 70 µl of T-PER Tissue Protein Extraction Reagent (ThermoScientific, Rockford, IL, USA). After incubating for 20 min on ice, samples were centrifuged for 10 min at 16,000 *x g* and the supernatant was transferred into a new microcentrifuge tube. Protein concentrations were measured using the Pierce BCA Protein Assay Kit (ThermoScientific, Waltham, MA, USA). Total protein lysates (15-50 µg) were subjected to SDS-PAGE using NuPAGE 4-12% Bis-Tris Midi Protein Gels (Invitrogen, Carlbad, MA, USA) and subsequently transferred to Polyvinylidene difluoride (PVDF) membranes using the iBlot 2 Dry Blotting System (Invitrogen, Carlbad, MA, USA). Unspecific antibody binding was prevented by incubating the blots in blocking solution containing 5% non-fat milk or 5% BSA (Sigma-Aldrich, St. Louis, MO, USA) in 1x Tris-buffered saline with 0.1 % Tween 20 (TBST) for 1 h at RT. Proteins were detected from the blots using the following primary antibodies diluted in the appropriate ratio with 1x TBST and incubated overnight at +4°C: rabbit anti-phospho-Akt1 (S473, 1:1000, #9018, Cell Signaling Technology, Danvers, MA, USA), rabbit anti-phospho-Akt2 (S474, 1:1000, #8599, Cell Signaling Technology, Danvers, MA, USA), rabbit anti-phospho-Akt (Thr308/309/305, 1:1000, #13038, Cell Signaling Technology, Danvers, MA, USA), rabbit anti-Akt1 (1:1000, #75692, Cell Signaling Technology, Danvers, MA, USA), rabbit anti-Akt2 (1:1000, #3063, Cell Signaling Technology, Danvers, MA, USA), rabbit anti-Akt (1:1000, #9272, Cell Signaling Technology, Danvers, MA, USA), rabbit anti-phospho-GSK3β (S9, 1:1000, #9336, Cell Signaling Technology, Danvers, MA, USA), rabbit anti-GSK3β (1:1000, #9315, Cell Signaling Technology, Danvers, MA, USA), custom-made mouse anti-phospho-Tau (B6, 1:1000) [17], mouse anti-4R-Tau (RD4, 1:1000, 05-804, Millipore), mouse anti-SQSTM1/p62 (1:1000, #5114, Cell Signaling Technology, Danvers, MA, USA), mouse anti-LC3 (1:1000, ab51520, Abcam, Cambridge, UK), mouse anti-Caspase-3 (1:1000, #9662, Cell Signaling Technology, Danvers, MA, USA), mouse anti-phospho-ERK (1:500, sc-7383, Santa Cruz Biotechnology, Dallas, TX, USA), rabbit anti-ERK2 (1:500, sc-154, Santa Cruz Biotechnology, Dallas, TX, USA), mouse anti-p85α (1:1000, ABIN1098111, Antibodies-online GmbH, Aachen, Germany), rabbit anti-phospho-SYK (1:1000, MA5-14918, Invitrogen, Waltham, MA, USA), rabbit anti-SYK (1:1000, #13198, Cell Signaling Technology, Danvers, MA, USA), mouse anti-β-actin (1:1000, ab8226, Abcam, Cambridge, UK) and mouse anti-GAPDH (1:15000, ab8245, Abcam). Blots were subsequently probed with the appropriate horseradish peroxidase (HRP)-conjugated secondary antibodies, either sheep anti-mouse-HRP (1:5000, NA931V, GE Healthcare, Chicago, IL, USA) or donkey anti-rabbit-HRP (1:5000, NA934V, GE Healthcare, Chicago, IL, USA) diluted in 1x TBST and incubated for 1 h at room temperature. Enhanced chemiluminescence (ECL, GE Healthcare, Chicago, IL, USA) was used to detect the protein bands. Blots were imaged with the Chemidoc MP system (Bio-Rad, Hercules, CA, USA) and images were quantified using the ImageLab (Bio-Rad, Hercules, CA, USA) software.

### AlphaLISA assay

Phospho-Syk level from WT mouse primary microglia cultures were determined using AlphaLISA SureFire Ultra kit (PerkinElmer, Waltham, MA, USA) according to manufacturer’s instructions. Briefly, after treatments, cells were lysed in freshly prepared 1X Lysis buffer (40µl/well) and agitate on a plate shaker (~350 rpm) for 10 minutes at room temperature. 30 μl of the lysate was transferred to a 96-well 1/2AreaPlate™ for assay. Acceptor Mix was added (15 µl/well) and plate was sealed with Topseal-A adhesive film. Plate was incubated for 1 hour at room temperature covered with foil. Donor Mix was added to wells (15µl/well) under subdued light. Plate was sealed with Topseal-A adhesive film, covered with foil, and incubated for 1 hour at room temperature in the dark. Plate was analyzed with EnVision plate reader (PerkinElmer, Waltham, MA, USA), using standard AlphaLISA settings.

### ELISA and Nitric Oxide assays

Aβ40 and Aβ42 levels in the mouse hippocampus homogenates were determined with monoclonal and HRP-conjugated antibody-based Human/Rat β amyloid 40 ELISA kit (Wako, Osaka, Japan). Mouse TNF-α and IL-6 ELISA Ready-SET-Go! kits (Affymetrix, San Diego, CA, USA) were used for the detection of tumor necrosis factor α (TNF-α) and IL-6 in the conditioned media of WT and Akt2 KO mouse primary microglia cultures treated with LPS (200ng/ml, Sigma-Aldrich, St. Louis, MO, USA) and IFNγ (20ng/ml, Sigma-Aldrich, St. Louis, MO, USA) for 24h. Nitric oxide (NO) levels were determined using the Griess Reagent Kit for Nitrite Determination (G-7921, Life Technologies, Eugene, OR, USA). All kits were used as recommended by the manufacturers.

### RNA sequencing

Library preparation and sequencing was conducted in Finnish Functional Genomics Centre (FFGC), Turku Centre for Biotechnology, University of Turku and Åbo Akademi University. The quality of the total RNA samples as well as prepared libraries was ensured with Advanced Analytical Fragment Analyzer. Sample and library concentration were measured with Qubit® Fluorometric Quantitation, Life Technologies. Library preparation was done according to Illumina TruSeq® Stranded mRNA Sample Preparation Guide (part # 15031047). The 45 libraries with good quality were pooled in one pool and run in 2 lanes. The samples were sequenced using Illumina HiSeq 3000 instrument to a minimum read depth of 12 million reads per sample. Single-read sequencing with 1 x 50 bp read length was used, followed by 8 + 8 bp dual index run.

### Quality control and transcript abundance quantification

FastQC version 0.11.8 (http://www.bioinformatics.babraham.ac.uk/projects/fastqc/) was used to examine the quality of the RNA sequencing reads. Illumina sequencing adapter sequences were removed, and reads were quality trimmed using the Trimmomatic version 0.38 [7]. The obtained trimmed reads were then mapped against the ribosomal and mitochondrial reference sequences (build mm10) using the Bowtie 2 version 2.2.3 [52]. Successfully mapped reads were abandoned. The rest of the reads were subjected to pseudo alignment to mouse reference transcriptome (build mm10) and transcript abundance quantification using kallisto version 0.44.0 [8].

### Differential expression analysis

Transcript abundance estimates were collapsed to gene-level counts using R package tximport version 1.10.1 [90]. Gene-level counts were pre-filtered (mean of gene count > 10), normalized and variance stabilizing transformed using R package DESeq2 version 1.22.2 [56]. Potentially confounding batch effects were corrected using removeBatchEffect function from limma version 3.38.3 [79]. The differentially expressed genes between diet and genotype groups were analyzed using the DESeq2.

### Consensus weighted gene co-expression network analysis

We carried out Consensus Weighted Gene Co-expression Network Analysis (WGCNA) as described previously [51]. Since the experimental design contains two variables of interest (genotype and diet), we carried out a consensus network analysis of two datasets: data from AwTw, AwT+, A+Tw, and A+T+ mice on STD, and data from the same genotypes with TWD. The rationale is that a consensus analysis identifies modules that group together genes correlated in both datasets, i.e., both with respect to diet as well as genotype. A thresholding power of 6 was chosen as it was the smallest threshold that resulted in a scale-free R^2^ fit of 0.8). The “signed hybrid” network was used (deepSplit=4, minModulesize=40) in which negatively correlated genes are considered unconnected. This analysis identified 28 co-expression modules ranging from 52 to 2728 genes per module (Supplementary Table 1). Genes in each module were further represented by a single representative expression profile (module eigengene, 1^st^ principal component of the module). Module eigengenes were correlated with genotype to determine the association of co-expressed genes with genotype. Additionally, module eigengenes allow one to define a continuous measure of membership of all genes in all modules. Genes with high module membership in a module are called hub genes for the module. To perform differential analysis of the consensus module networks (STD and TWD ~ genotype networks) we correlated and hierarchically sorted the identified module eigengenes for each network. Evaluation of module connectivity conservation was done by comparison of module sorting in their respective dendrograms.

### Gene set enrichment analysis

We used the R package anRichment (https://horvath.genetics.ucla.edu/html/CoexpressionNetwork/GeneAnnotation/) to calculate the enrichment of co-expression gene modules in a collection of reference gene sets that includes Gene Ontology terms, KEGG pathways, literature gene sets collected in the userListEnrichment R function [61], Molecular Signatures Database gene sets [92], aging gene sets from Enrichr [11], microglia-relevant gene sets from several recent articles [10, 18, 19, 40, 49, 100] and other gene sets. Fisher’s exact test was used to evaluate overlap significance.

### Transcription factor enrichment analysis

Over-represented transcription factor binding motifs (TFBS) and potential transcription factors (TF) in the surroundings of the transcription start site (TSS) of the differentially expressed genes (adjusted p-value <0.005 due to diet) were identified using R package RcisTarget version 1.4.0 [1]. We used genome wide TFBS ranking database containing regions from 500bp_upstream_and_100bp_downstream from TSS (motif collection mc9nr / mm10). Identified significantly enriched TFBS (normalized enrichment score (NES) threshold > 4.0) were annotated to TFs by using provided annotation database.

### Cell type enrichment analysis

Cell type enrichment was determined by cross-referencing of module genes with lists of genes known to be preferentially expressed in different cell types in mouse brain [86, 107]. All identified genes were used as background. Significance of enrichment of cell type markers with clusters was assessed using one tailed Fisher’s exact test.

### Statistics

IBM SPSS version 25 and R were used to analyze the data. Kolmogorov-Smirnov test was used first to test the normality of the distributions. The body weight at 12 months of age, blood glucose levels in the glucose tolerance test, spontaneous activity, latency to enter on day 2 in passive avoidance and search bias in the Morris swim task were analyzed with two-way ANOVA with A genotype, T genotype, and diet as between-subject factors. The task acquisition across days in Morris swim task was assessed with a mixed ANOVA for repeated measures (ANOVA-RM) using day as the within-subject and A and T genotypes as between-subject factors. β-Amyloid plaque load and the number of dystrophic neurites around plaques were analyzed separately for LEC and HC among A+ transgenic mice with two-way ANOVA using the T genotype and diet as factors. Statistical comparisons of biochemical analysis results were performed using two-way ANOVA followed by Fisher’s Least Significant Difference (LSD) post-hoc test. Statistical comparisons of correlations were performed using Spearman’s rho test. Results are expressed as mean ± standard error of mean (SEM) of control samples. P-values < 0.05 were considered statistically significant.

## Results

### Feeding of mice with TWD results in T2D

At the beginning of the dietary intervention, mice in the different test groups weighed on average between 22.5 – 24.4 g. There was no difference between APPswe/PS1dE9 genotypes (Aw vs. A+, p = 0.32), or Tau P301L genotypes (Tw vs. T+, p = 0.57) but mice in the to-be STD groups tended to weigh more than mice in the to-be TWD groups, p = 0.04). During the 6-month dietary intervention, there was a significant diet effect on the body weight increase (F_6,33_ = 25.6, p < 0.001), while the weight gain was less robust in A+ mice on TWD as compared to Aw mice (p = 0.01) (Fig. 1a). In the GTT, baseline glucose levels were higher (F_1,38_ = 3.8, p = 0.06; Fig. 1b) and 30 min glucose levels robustly elevated (F_1,38_ = 13.9, p = 0.001; Fig. 1c) in TWD groups as compared to STD groups. A possible fatty liver change was assessed in hematoxylin-eosin stained liver sections and blindly scored by two raters. Evidence of lipid vacuoles was found in 20/24 mice (69%) fed with TWD, but only in 5/22 mice (23%) fed with STD. Additionally, clear fatty liver changes were found in 12/24 TWD mice (50%), and in none of the STD mice (Fig. 1d). The diet effect was highly significant on the fatty liver score (p < 0.001, Mann-Whitney), while neither A (p = 0.32) nor T (p = 0.19) genotype influenced the score. In summary, feeding the mice with TWD results in a phenotype related to T2D irrespective of genotype.

### TWD exacerbates memory impairment associated with β-amyloid pathology

Previous studies have demonstrated that transgenic mice overexpressing the APPswe/PS1dE9 mutation are hyperactive [39, 80]. Here, A+ mice traversed a significantly longer distance during the 10-min test time than Aw mice (F_1,38_ = 15.7, p < 0.001), whereas the T genotype (p = 0.43) or diet (p = 0.84) did not influence spontaneous locomotion (Fig. 1e). Thus, to rule out the possible confounding effect of hyperactivity associated with the A+ genotype, we chose spatial navigation, where hyperactivity may speed up learning, and passive avoidance, where hyperactivity is unfavorable for the outcome, as tests for assessing possible memory impairment. In passive avoidance, all three factors (diet, A+, and T+ genotypes) showed a significant effect on the latency to enter the punished dark compartment, 48 h after the learning episode. A+ mice performed worse as compared to Aw mice (F_1,38_ = 8.8, p = 0.005), T+ mice worse than Tw mice (p = 0.02) and TWD mice worse as compared to STD mice (p = 0.02) (Fig. 1f). In the acquisition phase of Morris swim navigation task, there was no difference between the groups in their swimming speed (for all main effects p > 0.60). A+ mice had significantly longer escape latencies than Aw mice (F_1,38_ = 11.6, p = 0.002), whereas the T genotype (p = 0.27) and diet (p = 0.46) did not influence escape latencies (Fig. 1g). In the probe test on day 5 for search bias, A+ mice spent less time in the former platform zone than Aw mice (F_1,38_ = 12.6, p = 0.001) while the T genotype did not affect the search bias (p = 0.64). TWD tended to further impair the search bias in the groups with mixed genotypes (AwT+ or A+Tw; Fig. 1h), but overall, the diet main effect was not significant (p = 0.10). Altogether, this suggests that TWD exacerbates memory impairment in mice associated with β-amyloid pathology.

### TWD-induced T2D does not influence β-amyloid plaque load but increases the number of neuritic plaques

An obvious question behind the exacerbation of memory impairment of A+ mice by TWD is whether the diet aggravated the brain β-amyloid pathology. This question actually has two components; first, whether the dietary intervention affects the brain β-amyloid load, and second, whether it affects the formation of dystrophic neurites around the β-amyloid plaques, i.e. formation of neuritic plaques that are the most characteristic pathological feature of AD. To this end, we analyzed β-amyloid load in A+ mice in the lateral entorhinal cortex (LEC) and dentate gyrus of hippocampus (HC) (Fig. 1i), the brain sites with the earliest pathological changes in human AD and substantial β-amyloid plaque pathology in the APPswe/PS1dE9 mouse model. One mouse in the A+Tw STD group was removed as an outlier in this analysis to obtain normal distribution of data (Fig. 1j and k). Neither the T+ genotype (F_1,21_ = 0.4, p = 0.52) nor diet (p = 0.17) significantly influenced the β-amyloid load in LEC (Fig. 1 j) or in HC (Fig. 1k, T: p = 0.50, diet: p = 0.74). Accordingly, no significant effect by T+ genotype or diet on soluble Aβ42 (Supplementary Fig. 1a, T: F_1,17_ = 0.006, p=0.94, diet: F_1,17_ = 0.60, p=0.45), Aβ40 (Supplementary Fig. 1b, T: F_1,17_ = 1.0, p=0.32, diet: F_1,17_ = 3.4, p=0.08), or Aβ42/40 ratio (Supplementary Fig. 1c, T: F_1,17_ = 0.003, p=0.95, diet: F_1,17_ = 0.37, p=0.55) was found in hippocampal lysates of these mice. Next, we assessed the number of phospho-Tau (AT8)-positive dystrophic neurites around β-amyloid plaques (Fig. 1l). To this end, we counted the number of β-amyloid plaques above the threshold diameter of 9.9 µm in three LEC and three HC sections in each mouse, yielding 58-119 plaques per mouse in LEC and 47 – 84 plaques per mouse in HC. Recent evidence suggests that the number of dystrophic neurites around β-amyloid plaques in transgenic APP mice increases with the plaque diameter [5]. We found a similar relationship between the plaque diameter and the number of neurites both in LEC (R_1923_ = 0.28, p < 0.001) and HC (R_1467_ = 0.36, p < 0.001) (data not shown). Importantly, the average plaque diameter did not differ between the diet groups either in LEC (t_1921_ = 0.48, p = 0.63) or in HC (t_1465_ = 1.5, p = 0.14) (data not shown). However, TWD significantly increased the number of neurites around β-amyloid plaques in both LEC (F_1,1919_ = 80.3, p < 0.001) and HC (F_1,1463_ = 6.9, p = 0.009), while the T genotype had no effect in either LEC (p = 0.11) or in HC (p = 0.07)(Fig. 1m and n). Collectively, phenotypic and biochemical assessments show that TWD exacerbates memory impairment associated with β-amyloid pathology. However, TWD does not alter total β-amyloid load, β-amyloid peptide ratios or plaque diameter in mouse brain, but instead significantly increases the number of plaque-associated dystrophic neurites.

### TWD suppresses global transcriptional response to AD pathology in the hippocampus

To evaluate the impact of TWD on the AD-associated transcriptional response, we performed RNA-sequencing on hippocampal samples from AwTw, AwT+, A+Tw and A+T+ mice on STD or TWD. Assessment of the primary sources of variance in the dataset by principal component analysis (PCA) revealed gradual separation between AwTw, AwT+, A+Tw, and A+T+ mice, with A+T+ showing greatest separation from AwTw mice, suggesting that the combination of β-amyloid and Tau pathologies generates the most potent pathological state (Supplementary Fig. 2b). Separate PCA for mice of each genotype (to dissect diet effects) showed defined separation between STD and TWD samples in all genotypes, suggesting an effect for TWD on the transcriptional landscape of the hippocampus upon both β-amyloid and Tau pathology, as well as in the healthy state (Supplementary Fig. 2c). Surprisingly, almost no genes were differentially expressed (DE; false discovery rate (FDR) < 0.05) in AwT+ mice on both STD (3 up, 1 down) or TWD (1 up, 0 down), when compared to respective AwTw mice (Supplementary Fig. 2d, Supplementary Table 1-2). In turn, we observed 203 DE (199 up, 4 down) genes in A+Tw/STD mice and 173 DE genes (166 up, 7 down) in A+Tw/TWD mice. In accordance with the notion that β-amyloid and Tau pathologies potentiate each other’s effects, we observed 338 DE genes (294 up, 44 down) in A+T+/STD mice, but only 177 DE genes (162 up, 15 down) in A+T+/TWD mice. In general, a smaller amount of DE genes was observed in TWD mice when compared to STD mice, which was most evident in A+T+ mice. Comparison of the DE genes between STD and TWD mice for each genotype show a large overlap in genes, with unique DE genes mainly observed in STD mice (Supplementary Fig. 2d), suggesting a similar but less potent response in TWD mice when compared to STD mice. To further compare the transcriptional response between STD and TWD mice, we correlated DE analysis Z-statistic values for STD and TWD mice for each genotype. A strong positive correlation can be considered a highly similar transcriptional response. This analysis revealed only modest correlations (AwT+, cor = −0.309 (data not shown); A+Tw, cor = 0.415; A+T+, cor = 0.165, Fig. 2a), supporting the notion that TWD mice display an impaired transcriptional response to AD-related pathologies. Next, we performed enrichment analyses of DE genes (Supplementary Table 3) using a curated collection of both public gene sets (e.g., GO and MSigDB) and self-curated gene set collections (Methods). Consistent for both A+Tw and A+T+ mice, the top enrichment term for upregulated genes was “Top human microglia-specific genes”, irrespective of diet. Interestingly, a significant enrichment for synaptic function-associated terms (e.g. “chemical synaptic transmission” p = 6.7×10^−03^) was only observed for genes downregulated in A+T+/TWD mice. Enrichment analysis of genes showing discordant response in mice with A+ or A+T+ background upon TWD as compared to STD mice (Fig. 2a, blue points: Z > 2.5 or < −2.5 in STD, but not in TWD) indicated enrichment for microglia-specific terms (e.g. “Top human microglia-specific genes”, “Genes correlated with Trem2”), suggesting that TWD largely alters the transcriptional response of microglia (Supplementary Table 4). Expression profiles of genes displaying a discordant response revealed that the majority of genes were either up- or downregulated already in AwTw mice, showing only minor changes in relation to β-amyloid and/or Tau pathology. This could suggest that TWD locks microglia to a type of intermediate state (Supplementary Fig. 3a). Taken together, these results suggest a large range of transcriptional changes due to TWD, which potentially alter the response of microglia and neurons to β-amyloid and/or Tau pathology as compared to STD mice.

**Fig. 2.**
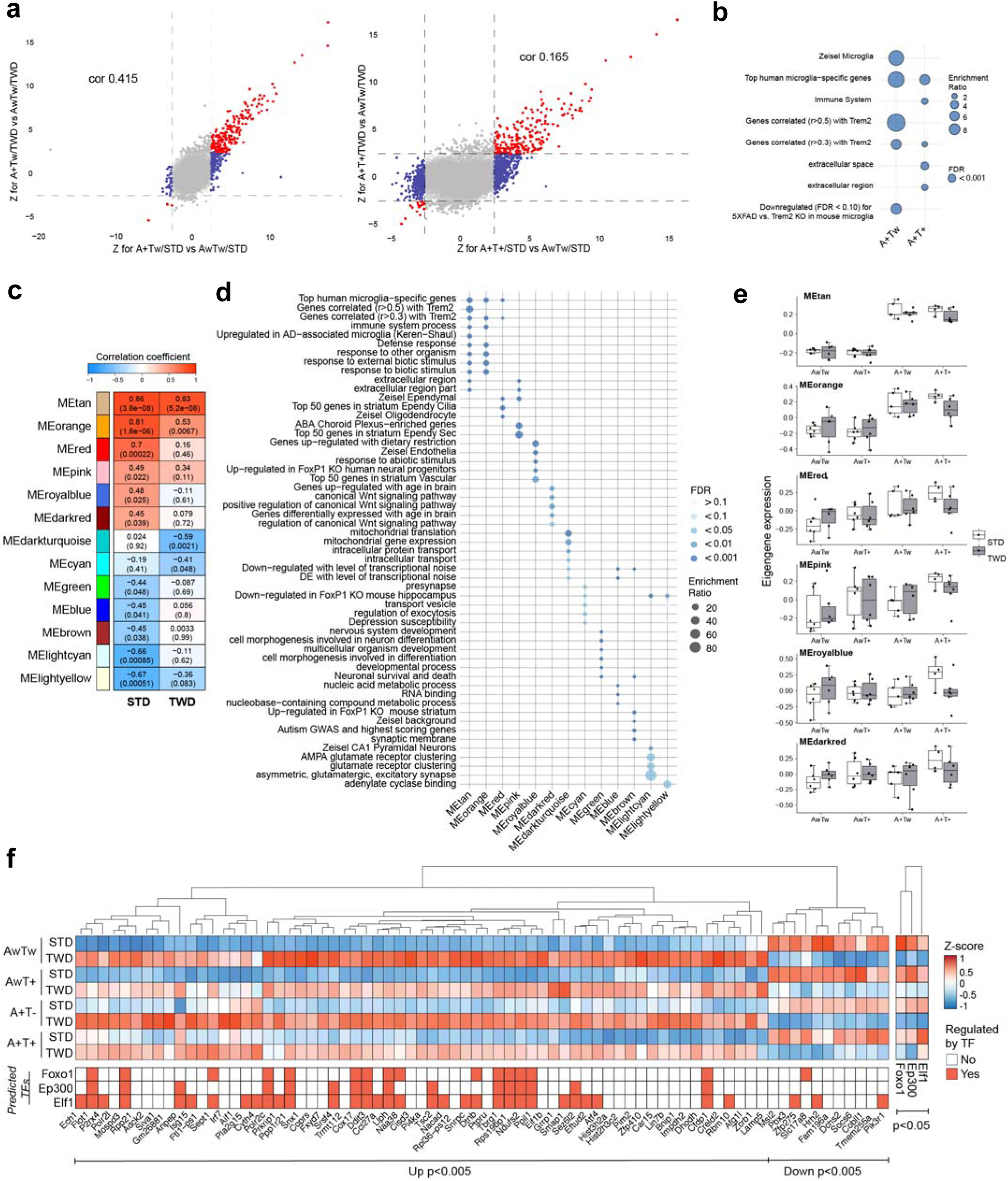
Transcriptomic and co-expression network analysis reveal impaired microglial response to AD-related pathology. **a** Transcriptome-wide response comparison of STD and TWD mice to A+ and A+T+ background. Plots show Z-statistics derived from DE analysis for A+Tw/STD vs. AwTw/STD (x-axis) and A+Tw/TWD vs. AwTw/TWD (y-axis) (left) and A+T+/STD vs. AwTw/STD (x-axis) and A+T+/TWD vs. AwTw/TWD (y-axis) (right). Points (genes) showing concordant response in STD and TWD cases are indicated in red, and discordant response for TWD as compared to STD, in blue. Genome-wide correlations of Z statistics and the corresponding correlation values are indicated on the top of each panel. **b** Enrichment analysis of genes showing discordant response to A+ or A+T+ background in TWD mice as compared to STD mice (blue dots in A). Enrichment ratio = number of observed divided by the number of expected genes. **c** WGCNA modules significantly associated to genotype in either STD or TWD mice. The color and value in the box represent correlation coefficient value. The value in parentheses represents corresponding correlation p-value. **d** Enrichment analysis for WGCNA module genes. Color indicates FDR value and size enrichment ratio. **e** Box plots representing eigengene values for modules positively associated with genotype. Points represent eigengene values for each mouse in that group. Box plots show the median, 25th and 75th percentiles, error bars show 1.5 interquartile ranges. **f** Z-score value heatmap of DE genes for STD vs. TWD mice with FDR < 0.005 and enriched transcription factors (TFs) with FDR < 0.05. n=4-6 mice/group.

### TWD diminishes β-amyloid and Tau pathology-associated microglial gene expression signatures

To further identify AD pathology-associated molecular networks that may be selectively affected by TWD in mouse brains, we performed consensus weighted gene co-expression network analyses (WGCNA; [51]). Our consensus WGCNA analysis identified 28 co-expression modules (Supplementary Table 5; Fig. 2c). By relating the module eigengene (a representative vector for the genes in a module, see methods), [51] to genotype, we identified six modules significantly positively associated and seven modules significantly negatively associated with genotype in either STD or TWD mice (Fig. 2c). A positive association can be considered as an increase in the expression of module genes in response to A+ and/or T+ background, and vice versa for negative association. A weaker association for the majority of the positively and negatively related modules in TWD mice was detected (Fig. 2c). Conversely, two negatively associated modules (MEdarkturquoise, MEcyan) showed a weaker association in STD mice (Fig. 2c). Comparison of the relations between STD and TWD network modules similarly revealed a loss in network preservation upon TWD as compared to the STD network, largely for positively associated modules, especially the MEdarkred module (Supplementary Fig. 3b).

Cell type enrichment analysis of the network modules revealed a significant enrichment for neuronal and astrocytic markers for negatively regulated modules, and for microglial and endothelial cell markers in positively correlated modules (Supplementary Fig. 3c). Accordingly, modules that negatively correlated with genotype were significantly enriched for neuron-related terms (Supplementary Table 6). In turn, top positively correlated modules were significantly enriched for “Top Human Microglia-Specific Genes”, and more specifically “Immune Response”, “Genes Correlated with Trem2”, and genes “Upregulated in AD-associated Microglia (Keren-Shaul et al., 2017)”. This suggests that the top identified positively associated modules recapitulate a microglial Trem2-driven DAM response, which is partially impaired upon TWD. MEdarkred, which had a poor preservation between the STD and TWD networks, was significantly enriched in genes related to the Wnt signaling pathway. Previously, it was shown that downregulation or knock-out of Trem2 in microglia results in reduced Akt S473 and Gsk3β S9 phosphorylation, resulting in β-catenin degradation and impaired Wnt/β-catenin [109]. These results suggest that TWD in mice potentially alters microglial functionality downstream of Trem2 signaling, thus resulting in the observed partial transcriptional response when compared to STD mice.

To assess whether any modules associated with Tau pathology, β-amyloid pathology or the combination of both, we plotted eigengene expression levels for each genotype and diet (Fig. 2e, Supplementary Fig. 3d). The most evident pathology-dependent alterations were observed in the top two positively associated modules, MEtan and MEorange, where an increase in eigengene expression was driven primarily by the presence of β-amyloid pathology. In turn, MEdarkturquoise, MEgreen, MElightyellow, MEblue, and MElightcyan, and MEred and MEpink modules showed a more linear change in expression (in order: AwTw, AwT+, A+Tw, A+T+), with the greatest decrease or increase in A+T+ mice when compared to AwTw mice. Interestingly, MEroyalblue and MEdarkred modules showed only modest or no changes in expression in AwT+ or A+Tw mice, but an increase in expression in A+T+ mice. This suggests that in some cases β-amyloid and Tau cooperate to cause specific transcriptional alterations in mouse brain, which is in line with recent findings [74]. Overall, our network analysis reveals a strong transcriptional response to β-amyloid pathology and combined β-amyloid and Tau pathologies in genes predominantly expressed in microglia and associated with microglia-specific biological terms. Specifically, a Trem2-driven DAM type profile was recapitulated, and this was partially impaired in TWD mice, potentially due to altered downstream signaling of Trem2.

### TWD induces aberrant expression of disease-associated microglia signature genes in mouse hippocampus

As TWD was found to change the transcriptional landscape in mouse hippocampus in WT (AwTw) and transgenic mice (AwT+, A+Tw, A+T+), resulting in an altered basal state and response to β-amyloid and Tau pathology, we next concentrated on the genes that were altered by the diet irrespective of genotype to pin-point potential key TWD-associated pathways. DE analysis for TWD mice vs. STD mice revealed 4428 DE genes (2698 up, 1730 down, Fig. 2f, Supplementary Table 7), enriched for terms related to ribosomes, mitochondria (e.g. “mitochondrial protein complex”), and genes up-/downregulated in AD-/ALS-associated microglia (“Up-regulated in AD-associated microglia [40], “Upregulated in Stage2 vs. Stage1 disease-associated microglia”) (Supplementary Table 8). Given that our genotype and diet DE analysis and network analysis suggested that TWD alters the previously described DAM response, we examined the overlap between TWD DE genes and DAM genes ([40], genes with FDR < 0.01 and FC > 3 or < −3). This analysis revealed an overlap of 132 genes with top DE hits, such as *Pik3r1* (FDR = 0.004, Log2FC = −0.40), *Tyrobp* (FDR = 0.006, Log2FC = 0.37), and *Tmem119* (FDR = 0.008, Log2FC = 0.29). The expressional changes of *Pik3r1*, *Trem2*, and *Tyrobp* detected by RNA-sequencing were confirmed using conventional qPCR (Supplementary Fig. 4). Despite observing a general lack of transcriptional response associated with Trem2 correlated genes, we observed an increase in both *Trem2* and *Tyrobp* expression levels. However, the expression of *Pik3r1*, a gene encoding PI3K regulatory subunit 1 and directly acting downstream of *Trem2*, was decreased. This finding highlights *Pik3r1* as a potentially important candidate in conveying the observed TWD effects, given the importance of the PI3K-mediated insulin signaling in the brain, the role of the Trem2-PI3K-Akt pathway in DAM activation, and the observed alterations in Trem2- and Wnt signaling pathways in our WGCNA analysis. Lastly, as TWD causes a large scale of transcriptomic effects, we sought to determine the transcription factors potentially regulating the observed top DE genes (p<0.005). Enrichment for three significantly differentially expressed transcription factors, *Foxo1*, *Ep300*, and *Elf1* was identified (Fig. 2f).

### TWD leads to decreased Akt and increased Gsk3β activation in hippocampus

Since analysis of the RNA-sequencing data revealed decreased expression levels of *Pik3r1*, the gene encoding p85α subunit of PI3K, in hippocampus of mice with TWD as compared to STD, we next assessed whether there are alterations in well-known downstream targets in PI3K-Akt-Gsk3β-Tau pathway. We analyzed hippocampal protein lysates using Western blot and found that phosphorylation of S473 in Akt1 kinase, a residue regulating Akt1 activation, was significantly decreased in TWD mice as compared to STD mice (F_1,38_ = 30, p<0.001, Fig. 3a). This was an expected finding, considering the decreased p85α expression. In line with this, phosphorylation of the corresponding S474 site in Akt2 (F_1,38_ = 9.9, p=0.003, Fig. 3b) and of another activating site, T308/309/305 in Akt1/2/3, respectively, (F_1,38_ = 28.9 p<0.001, Fig. 3c) were significantly decreased in the hippocampus of TWD mice as compared to STD mice. Furthermore, Akt-mediated phosphorylation of the inhibitory S9 residue in Gsk3β, a well-characterized Tau-kinase, was significantly decreased in mice on TWD as compared to STD (F_1,38_ = 14.3, p<0.001, Fig. 3d), suggesting increased Gsk3β activity. However, no diet effect was observed on Tau phosphorylation between the TWD and STD mice (F_1,37_ = 0.67, p=0.42), while Tau phosphorylation was significantly reduced in mice with T+ genotype (Supplementary Fig. 1d, F_1,37_ = 48, p<0.001). The typical 64-kDa band [82] present above the endogenous mouse Tau bands in T+ mice (P301L overexpressing human Tau) was included into the quantification. However, excluding the 64-kDa band from the analysis does not change the result. As expected, total Tau levels were significantly increased in T+ as compared Tw mice (Supplementary Fig. 1d, F_1,41_ = 141, p<0.001). The diet also had a significant effect on total Tau levels (F_1,41_ = 10.8, p=0.002). As previously described, hyperphosphorylated, aggregated Tau becomes insoluble, which might explain why increased Tau phosphorylation was not detected in hippocampal homogenates in this analysis. In contrast, IHC analysis showed that TWD significantly exacerbated dystrophic neurite i.e. phosphorylated Tau pathology in both A+Tw and A+T+ mice in the vicinity of β-amyloid plaques (HC and LEC, Fig 1l-n).

**Fig. 3.**
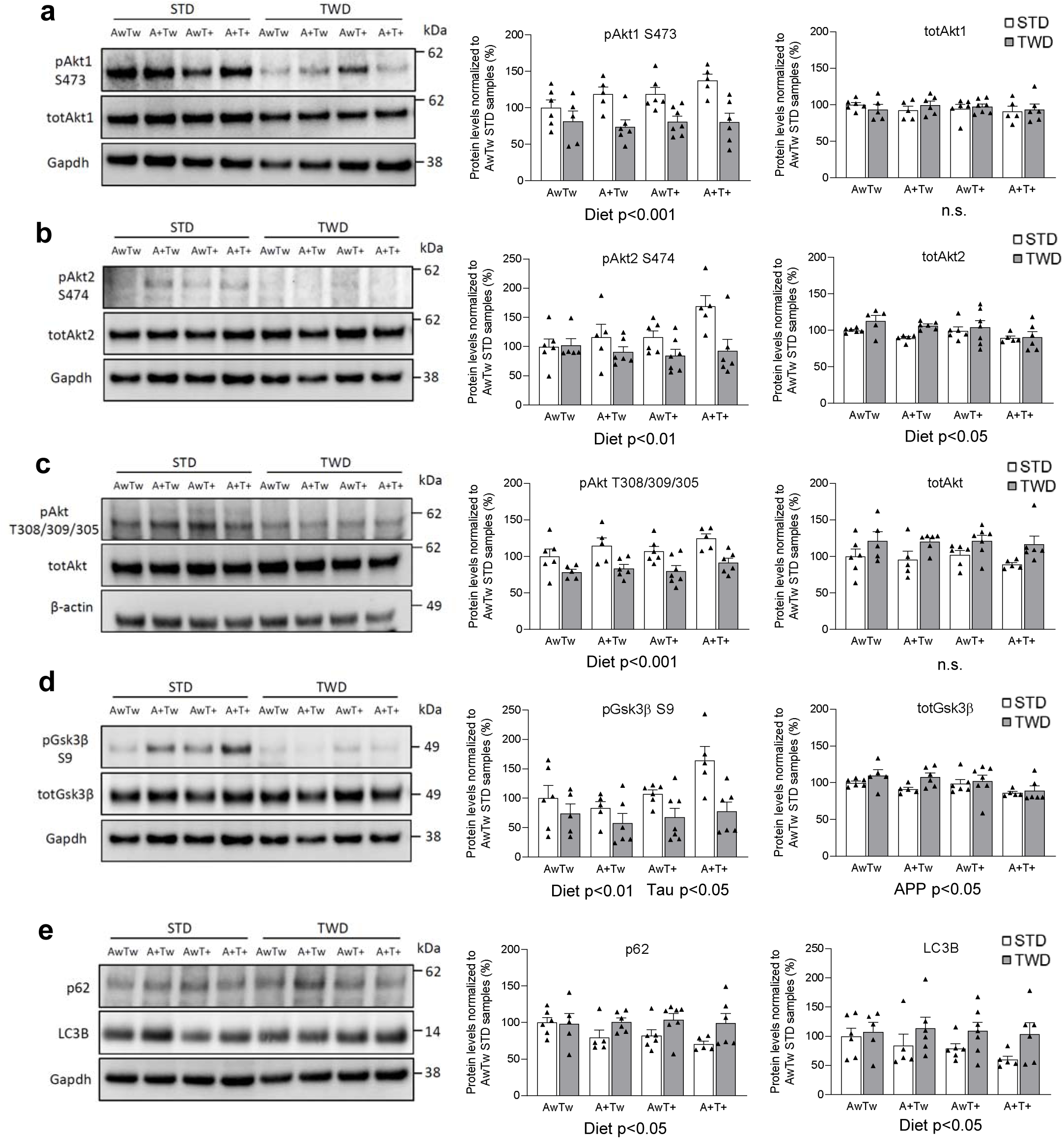
TWD induces impaired Akt-Gsk3β signaling in the brain. **a** Western blot analysis of hippocampal lysates showing decreased phosphorylation of S473 in Akt1 in mice with TWD as compared to STD (p<0.001). Diet had no effect on total Akt1 levels. **b** Similarly, phosphorylation of S474 in Akt2 is decreased in mice with TWD as compared to STD (p<0.01). Diet had a minor, but statistically significant effect on total Akt2 levels (p<0.05). **c** Also, phosphorylation of T308/309/305 residue in Akt1/2/3, respectively, was significantly decreased in mice with TWD as compared to STD (p<0.001). Total Akt1/2/3 levels showed no significant changes. **d** Phosphorylation of Gsk3β at the inhibitory S9 residue was significantly decreased in mice with TWD as compared to mice with STD (p<0.01). A+ genotype increased slightly, but significantly Gsk3β S9 phosphorylation (p<0.05) as well as total Gsk3β levels (p<0.05). **e** A representative Western blot image and quantification of hippocampal lysates showing increased levels of the autophagy markers p62 and LC3B in TWD mice as compared to STD mice (p<0.05). Phosphorylated protein levels were normalized to their respective total protein levels in cell lysates and total protein levels were normalized to Gapdh or β-actin. All results are shown as mean + SEM, n=5-7 mice/group, Two-way ANOVA.

Since energy metabolism and PI3K-Akt signaling are known to regulate autophagy, protein levels of well-known autophagy markers p62 and LC3B were next assessed in hippocampal lysates. Levels of p62 were significantly increased in TWD mice as compared to STD mice (Fig. 3e, F_1,38_ = 6.9, p=0.012), suggesting that TWD decreases autophagy. This was unexpected, since impaired activation of PI3K-Akt pathway has been shown to lead to decreased levels of p62 [46, 105] reflecting enhanced autophagic degradation. LC3B detection revealed increased levels of cytosolic LC3BI form and no membrane-bound lipidated LC3BII form in TWD mice as compared to STD mice (Fig. 3e, F_1,38_ = 6.3, p=0.016), further suggesting decreased autophagy.

Hippocampus is in a central role in processes related to learning and memory. Thus, we next analyzed whether the results of the behavioral tests correlated with any biochemical assessments in the hippocampal lysates. Higher body weight at age of 12 months (W12) and higher weight gain (WG) between 5 and 12 months negatively correlated with phosphorylation of Akt1 S473 (W12: R=-0.417, p<0.01 and WG: R=-0.453, p<0.01), Akt2 S474 (W12: R=-0.373, p<0.01 and WG: R=-0.373, p<0.01) and Gsk3β S9 (W12: R=-0.292, p<0.05 and WG: R=-0.331, p<0.05). However, no correlation between peripheral glucose intolerance and Akt/Gsk3β phosphorylation status was found. Interestingly, mean latency time to find the hidden platform in the acquisition phase of Morris swim task revealed a strong negative correlation with phosphorylation of Akt1 S473 (R=-0.355, p<0.016), Akt2 S474 (R=-0286, p=0.054), and Gsk3β S9 (R=-0.353, p<0.05), suggesting that the stronger the Akt/Gsk3β signaling, the faster the mice find the platform. However, spontaneous movement activity did not correlate with Akt or Gsk3β phosphorylation. Altogether, these results strongly suggest that decreased *Pik3r1* expression upon TWD coincides with significantly decreased activity of Akt1/2 (both S473/474 and T308/309/305 catalytic sites) and increased activity of Gsk3β. Furthermore, TWD exacerbates dystrophic neurite pathology in both A+Tw and A+T+ mice in the vicinity of β-amyloid plaques (HC and LEC, Fig 1l-n).

### TWD reduces clustering of microglia around β-amyloid plaques in mice

Since RNA-sequencing data also suggested that TWD leads to altered Trem2-signaling and general transcriptional response of microglia, we next assessed if TWD affected microglia-related pathology around β-amyloid plaques in mouse and human IHC samples. Hippocampal sections of A+Tw and A+T+ mice were triple-stained with anti-Iba1 (microglia), anti-Cd68 (lysosomal marker expressed in activated microglia) and X-34 (β-amyloid plaque) to assess microglia clustering and lysosomal activity around plaques. Iba1-positive area within 30 µm from the plaque was found to decrease in TWD as compared to STD mice (Fig. 4a-b, diet p<0.05). No diet effect on Cd68-positive area was observed. However, significant reduction of both Iba1 and Cd68 intensities were detected in mice fed with TWD as compared to mice fed with STD (Fig. 4c, diet p<0.001), suggesting that the number of microglia and lysosomal activity around plaques is reduced. Also, T+ genotype, on top of the A+ background, significantly increased both Iba1 and Cd68 intensity (Fig. 4c, p<0.001), suggesting that Tau pathology further potentiates microglia activation in A+ mice.

**Fig. 4.**
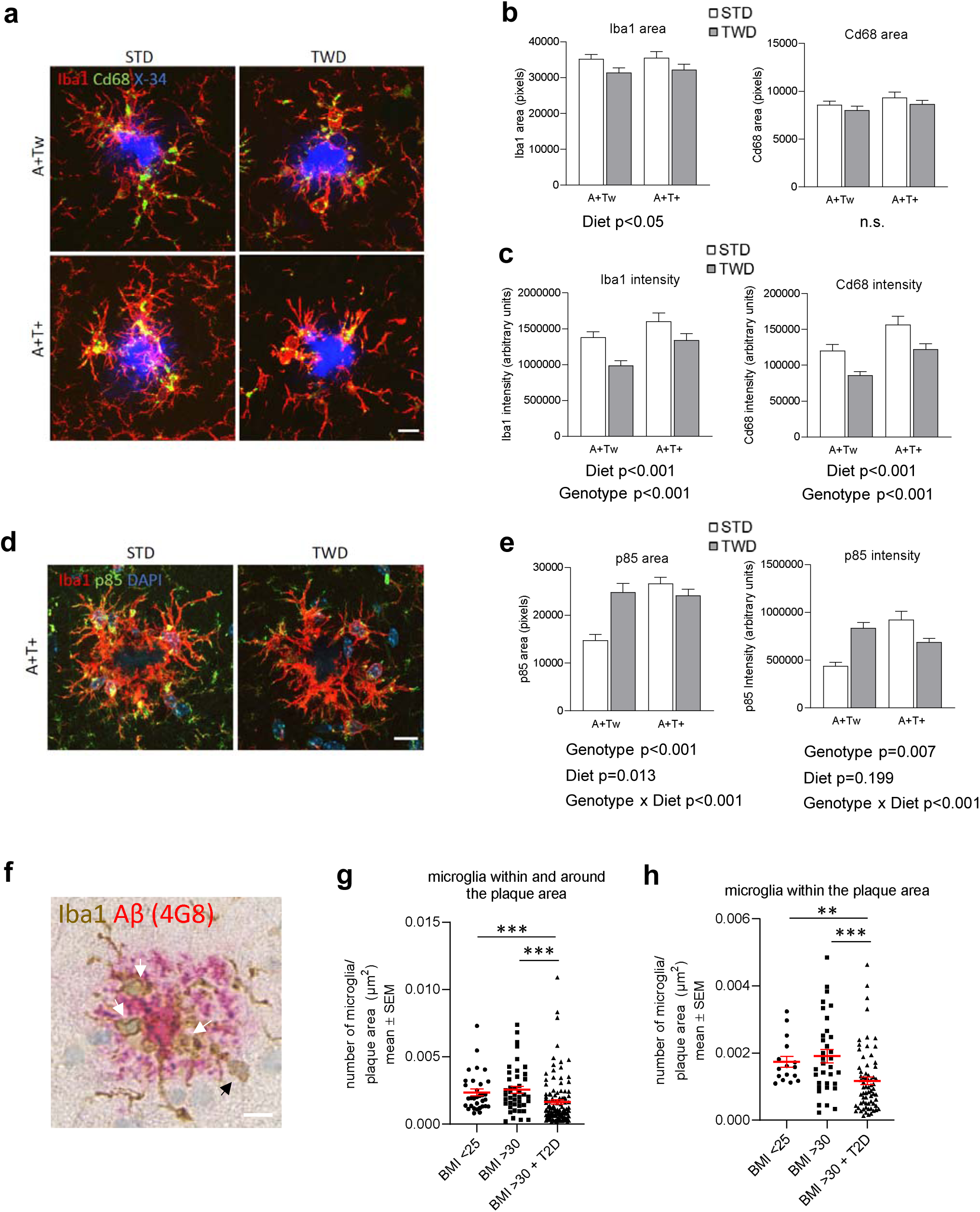
TWD and T2D alter microglial response to β-amyloid plaques in AD mice and human brain biopsies. **a** Representative immunofluorescence images of β-amyloid plaques stained with X-34 (blue) surrounded by Iba1 positive microglia (red) and lysosomes stained with Cd68 (green) from A+Tw and A+T+ mice from both STD and TWD groups. Scale bar 10µm. **b** Quantification revealed that TWD significantly (p<0.05) decreased Iba1 positive area (pixels) as compared to STD in both A+Tw and A+T+ mice, while diet showed no effect on Cd68 positive area. **c** Also, intensity of both Iba1 and Cd68 (arbitrary units) was significantly decreased (p<0.001, both) in mice with TWD as compared to STD in both genotypes. Furthermore, T+ genotype significantly increased both Iba1 and Cd68 intensity (p<0.001, both). **d** Representative immunofluorescence images of Iba1 positive microglia (red) and p85α staining (green) around β-amyloid plaques in hippocampus of A+T+ mice from both STD and TWD groups. Scale bar 10µm. **e** Quantification of p85α revealed statistically significant genotype x diet interaction (p<0.001) in both p85α area and p85α intensity. T+ genotype significantly increased both p85α area (p<0.001) and p85α intensity (p=0.007), while diet had opposing effects on p85α depending on the genotype. A-E; results are shown as mean + SEM, n=5-6 mice/group, Two-Way ANOVA. **f** Representative image of frontal cortex biopsy sample from probable iNPH subject co-stained for IBA1 (brown) and β-amyloid (red). White arrows denote microglia within the plaque area, and black arrow denotes microglia around the plaque area. Scale bar 10µm. **g** Quantification of IBA1-positive microglia within and around the β-amyloid plaque. **h** Quantification of IBA1-positive microglia within the plaque area. **g-h**; results are shown as mean ± SEM, n=3-5 subjects/group

Next, we wanted to assess whether the decrease of p85α expression observed at the transcriptional level could also be detected specifically in microglia around β-amyloid plaques in IHC samples. Hippocampal sections of mice with A+Tw and A+T+ genotypes were triple stained with anti-Iba1, anti-p85α and DAPI to assess p85α in microglial cells. Plaque-associated microglia were identified by the typical clustering pattern of Iba1 immunopositive microglia around the β-amyloid plaques (Fig. 4a and 4d). The analysis revealed a significant genotype x diet interaction on both p85α area and intensity (Fig. 4e, genotype x diet, F_1,247_ = 15.2, p<0.001). In A+Tw mice, TWD increased the levels of p85α in microglia surrounding the β-amyloid plaques. The T+ genotype also increased p85α levels in the microglia around β-amyloid plaques (Fig. 4e, T+ genotype, F_1,247_ = 15.9, p<0.001 for p85α area and F_1,247_ = 7.2, p=0.008 for p85α intensity) However, in contrast to A+Tw on TWD and A+T+ mice on STD, TWD induced a significant reduction in the levels of p85 in plaque-associated microglia in A+T+ mice. In conclusion, these data indicate that TWD reduces clustering of microglia around the β-amyloid plaques in A+Tw and A+T+ mice, which could be linked to the altered expression of p85α in the microglia.

### Metabolic phenotype affects microglia clustering around β-amyloid plaques in probable iNPH patients with β-amyloid pathology

To assess the effect of the metabolic phenotype on microglia in humans, we analyzed the extent of microglia clustering around β-amyloid plaques (Fig. 4f) in frontal cortical biopsies obtained from living patients with probable iNPH (Table 1). Subjects were divided in to three groups: normal weight individuals (BMI<25), obese individuals (BMI>30), and obese individuals with T2D diagnosis (BMI>30 + T2D). In accordance with our findings in TWD-fed mice, we observed a statistically significant decrease in the number of microglia around β-amyloid plaques in obese individuals with T2D as compared to both normal weight and obese individuals without T2D, when assessing microglia both around and within the plaque area (p<0.001) (Fig. 4g) as well as microglia only within the plaque area (p<0.001) (Fig. 4h). Interestingly, no difference was observed in the number of microglia between BMI<25 and BMI>30 groups. Thus, our results indicate that T2D, not obesity, reduces microglia clustering around β-amyloid plaques in the frontal cortical biopsies of probable iNPH patients.

### Modulation of PI3K-Akt signaling pathway affects phagocytosis and proinflammatory response in mouse microglia

Since TWD exacerbated dystrophic neurite pathology, reduced the clustering of microglia around β-amyloid plaques, and decreased the expression of *Pik3r1* in the hippocampus of A+Tw and A+T+ mice as compared to STD mice with the same genetic backgrounds, we next examined the microglia-specific molecular mechanisms underlying these observations. PI3K is a heterodimeric enzyme composed of the catalytic p110 subunit and the regulatory p85 subunit, which catalyzes the phosphorylation of PI(4,5)P_2_ to the lipid second messenger PI(3,4,5)P_3_ [41]. *Pik3r1* is abundantly expressed in mouse microglia [107] and it encodes three regulatory isoforms of PI3K (p85α, p55α and p50α), which are produced through alternative splicing [29]. To modulate PI3K-Akt signaling and to model the subsequent effects of decreased *Pik3r1* expression in mouse BV2 and neonatal primary microglia upon basal and LPS-induced stress conditions, we used the well-established PI3K inhibitor LY294002 and studied the effects on autophagosomal activity (the ratio of LC3BII/I), mitogen-activated protein kinase kinase (Mek)/extracellular signal-regulated kinase (Erk) pathway, apoptosis, and phagocytic uptake (Fig. 5a-f). The treatment of BV2 cells with LY294002 showed an expected dose-dependent decrease in the phosphorylation status of S473 in Akt (pAkt) in both normal and LPS-induced stress conditions (Fig. 5a and b), reflecting decreased Akt activity. LPS treatment alone significantly increased the levels of pAkt in BV2 cells as compared to untreated cells, which is consistent with our previous results [60]. In parallel with the reduced activity of Akt, the ratio of the autophagosomal marker LC3BII/I was increased in a dose-dependent manner as expected, particularly in the BV2 cells treated with both LPS and LY294002 (Fig. 5a-d and Supplementary Fig. 5). Conversely, the phosphorylation status of Erk1 and 2 (pErk1 and 2) in the activating Y204 site decreased in a dose-dependent manner in LY294002 samples, while the levels of active caspase-3 and p85α were unchanged in the BV2 cells treated with both LPS and LY294002 (Supplementary Fig. 5).

**Fig. 5.**
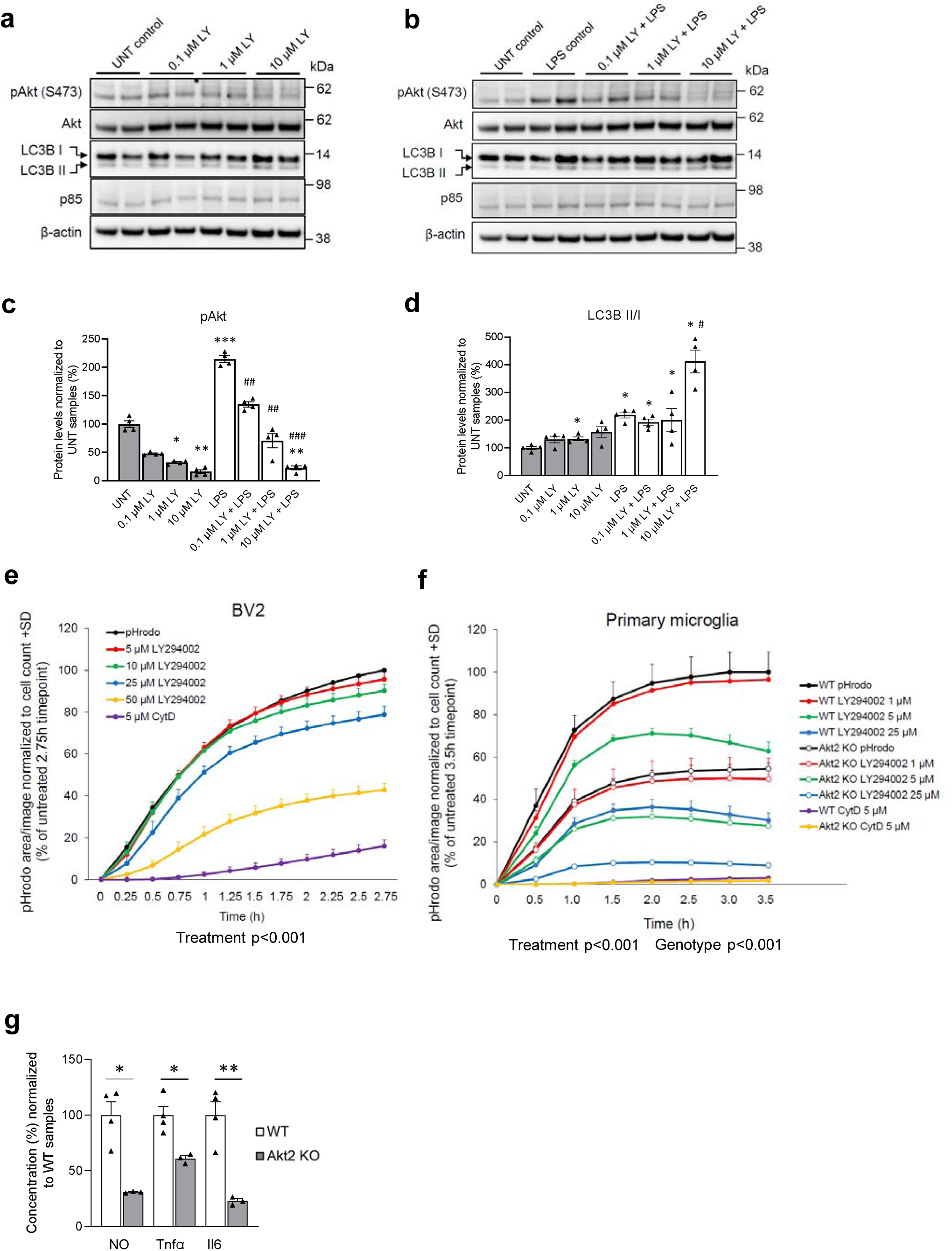
Modulation of PI3K-Akt pathway affects autophagic and phagocytic activity as well as inflammatory response in microglial cells. **a** A representative Western blot image of lysates of BV2 cells treated with LY294002 (LY) (0.1 µM – 10µM), and **b** with LY294002 together with LPS. **c** Quantification revealed that LY294002 decreased Akt S473 phosphorylation levels dose-dependently in cells treated with LY294002 and together with LPS and LY294002. **d** Quantification of LC3B II and I levels revealed a significant increase in LCB3II/I ratio by LY294002. **e** Phagocytic activity in BV2 cells was assessed using pHrodo bioparticles and fluorescence emission was measured in the IncuCyte live cell imaging device in every 15 min. BV2 cells were treated with four different concentrations of PI3K inhibitor LY294002 (5µM – 50µM) and with 5µM Cytochalasin D (CytD). LY294002 decreased phagocytic activity dose-dependently (p<0.001) and CytD blocked phagocytosis almost completely. a-d; Data are presented as mean + SEM from four biological replicates. Mann-Whitney U-test, *p<0.05, **p<0.01, ***p<0.001 vs. untreated control, #p<0.05, ##p#<0.01, ###p<0.001 vs. LPS control. **f** Phagocytic activity of primary microglia isolated from WT and Akt2 KO mice was assessed similarly as in BV2 cells using pHrodo bioparticles and fluorescence emission was measured in the IncuCyte live cell imaging device in every 30 min. Cells were treated with three different concentrations of LY294002 (1µM – 25µM) and with 5µM CytD. LY294002 decreased phagocytic activity dose-dependently (p<0.001) and Akt2 KO cells showed significantly decreased phagocytic activity as compared to WT cells (p<0.001). CytD blocked phagocytosis almost completely in both WT and Akt2 KO cells. e-f; Data are presented as mean + SEM from three biological replicates in technical quadruplets, One-way ANOVA, LSD (e) and Two-way ANOVA (f) **g** Levels of inflammatory markers, nitric oxide (NO), tumor necrosis factor α (Tnfα), and interleukin 6 (Il6) were significantly lower in Akt2 KO primary microglia as compared to WT primary microglia after 48h LPS+IFNγ -treatment. Data presented as mean % + SEM, n=3-4. *p<0.05, **p<0.01, T-test.

Next, live cell imaging of BV2 and primary microglia treated with LY294002 revealed a dose-dependent decrease in the phagocytic uptake of pHrodo-labeled bioparticles (Fig. 5e and f). Moreover, primary microglia isolated from neonatal Akt2 knock-out (Akt2 KO) mice also showed a significant decrease in the phagocytic uptake of pHrodo-labeled bioparticles as compared to WT microglia (Fig. 5f). The treatment of Akt2 KO microglia with LY294002 further decreased the phagocytic uptake of bioparticles as compared to untreated Akt2 KO microglia. Akt2 KO microglia cultures showed reduced levels of secreted Tnf-α, Il6 and nitric oxide (NO) in the culture medium as compared to the WT microglia cultures in response to LPS treatment (Fig. 5g). Altogether, these results suggest that the inhibition of PI3K-Akt signaling by using pharmacological (LY294002) or genetic (Akt2 KO mouse model) approaches significantly decreases the phagocytic uptake affects the autophagic and Erk kinase activity, and dampens the production of proinflammatory cytokines and NO upon LPS-induced stress in mouse microglia.

### Ligand-induced activation of Trem2/Dap12 upon the inhibition of PI3K-Akt signaling enhances the activation of spleen tyrosine kinase in mouse microglia

PI3K plays a central role in the Trem2/Dap12-mediated signaling in mouse macrophages and osteoclasts upon ligand-induced activation, resulting in alterations in Erk kinase activity, mobilization of calcium, reorganization of actin, and apoptosis [72]. Importantly, the same study showed that inhibition of PI3K prevented recruitment of p85α and spleen tyrosine kinase (Syk kinase) to Dap12. This is an intriguing finding given the fact that upon ligand binding to Trem2, the tyrosine residues within ITAM domain of Dap12 are phosphorylated, which leads to the recruitment of Syk kinase to activate downstream signaling molecules, such as PI3K, Erk, and phospholipase Cγ2 (Plcγ2) [72]. To assess whether inhibition of PI3K exerts a similar effects also in microglia, the phosphorylation of Dap12 and Syk kinase (Y525/526, pSyk) by Src-kinase was stimulated by using macrophage colony stimulating factor (M-CSF) in mouse BV2 microglia with or without LY294002 (Fig. 6a-c). M-CSF is known to activate Dap12 and Syk kinase through outside-in signaling within minutes after the treatment [72, 110]. The treatment of BV2 cells with M-CSF for five minutes resulted in the expected ~ two-fold increase in pSyk levels (p<0.001) as compared to control cells, which was abolished by Src-kinase inhibitor SU665 as shown previously [110](Fig. 6a and c). Unexpectedly, the combined treatment with LY294002 and M-CSF significantly increased the levels of pSyk in BV2 cells (p<0.05, Fig. 6a-c). To further pinpoint the observed inhibitory effect of PI3K on pSyk levels in the Trem2/Dap12 signaling pathway, we used Trem2 antibody for direct ligand-induced activation of Trem2 alone or in combination with LY294002 for five minutes (Fig. 6d). Alpha-LISA-based analysis of pSyk showed an average 2.5-fold increase in the levels of pSyk after the treatment of primary microglia cultures with Trem2 antibody as compared to control cultures (Fig. 6d). Similar to the combined treatment with LY294002 and M-CSF in the BV2 cells, co-treatment of primary microglia with LY294002 and Trem2 antibody significantly further potentiated the increased levels of pSyk. These results suggest that the inhibition of PI3K activity increases Syk kinase activity upon ligand-induced activation of Trem2/Dap12 signaling in the microglia.

**Fig. 6.**
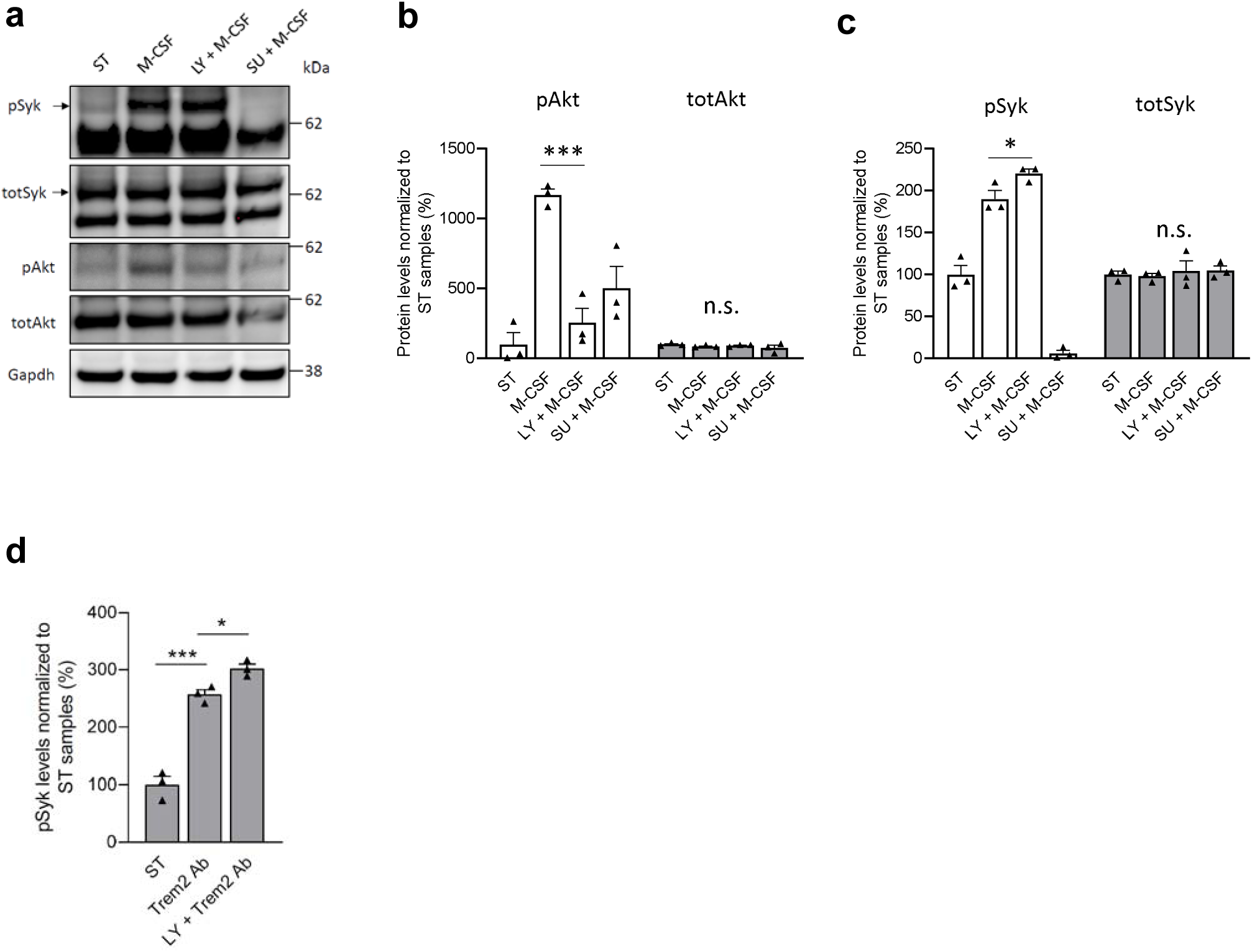
Inhibition of PI3K activity affects phosphorylation of Syk in microglial cells. **a** Western blot analysis of lysates of BV2 cells pre-treated with LY294002 (LY, 1µM) or Src-kinase inhibitor SU6656 (SU, 2µM) prior to M-CSF treatment. **b** Quantification of Akt S473 phosphorylation levels revealed a significant decrease in LY294002-treated samples while total Akt levels were unchanged. **c** Quantification of Syk (Y525/526) phosphorylation levels revealed a significant increase in LY294002-treated samples while total Syk levels were unchanged. **d** Phosphorylation status of Syk was significantly increased in WT primary microglia treated with Trem2 antibody for 5 min as compared to control cells (ST). LY294002 treatment together with Trem2 antibody further increased phosphorylation. ST; starvation only, M-CSF; 100ng/ml for 5 min, LY; 1µM LY294002 for 1 h before M-CSF, SU; 2µM Src-kinase inhibitor SU6656 treatment 20 min before M-CSF. Data presented as mean % + SEM, n=3, *p<0.05, **p<0.01, ***p<0.001, One-way ANOVA, LSD

## Discussion

TWD, which is manifested as increased intake of energy-dense foods high in saturated fat, sugar and cholesterol, and low in fiber, is a major contributor to the increasing prevalence of obesity and T2D, imposing a global health challenge. Several studies have shown that TWD, obesity, and T2D increase the risk of AD [14, 21, 31, 45, 50, 103]. Despite these seminal studies, the underlying molecular mechanisms remain elusive. Here, we have assessed the effects of six-month TWD exposure on memory, brain pathology, and global gene expression in female mice with different genetic backgrounds linked to AD and/or tauopathy (A+Tw, AwT+ and A+T+). In addition, we set the goal to address the translation of the mouse model-derived results to humans by utilizing cortical biopsies obtained from living iNPH patients presenting with β-amyloid pathology. In this unique human tissue sample set [53], we were able to specifically address T2D- and obesity-related effects in the frontal cortex of normal weight and obese iNPH patients with or without T2D. To our knowledge, this is the first study, which comprehensively addresses the full spectrum of AD-associated pathologies in parallel in several transgenic mouse lines and human cortical tissue samples with β-amyloid pathology to elucidate TWD-, T2D, and obesity-related alterations in cellular processes.

We successfully induced obesity and diabetic phenotype in all of the mouse lines used by applying a six-month TWD regime, starting at the age of seven months, which is in line with previous reports [26, 36, 73, 94]. Furthermore, TWD strongly increased fatty liver changes, emphasizing the robust effect of TWD on the metabolism of not only in the brain but also in peripheral tissues, as also shown previously [97]. Our results related to the behavioral assessments strongly suggest that TWD-induced obesity and diabetic phenotype impair memory and learning, which are also supported by the previous findings [24, 26, 33, 34, 38, 47, 81]. More specifically, the comparison of the four genotypes suggests that adverse effects of TWD on memory and learning are the most prominent in mice with a moderate genetic predisposition to develop AD-like brain pathology (A+Tw and AwT+), while in the A+T+ mice, the strong genetic burden overpowers the effect of TWD. Previous studies have shown that TWD and HFD associate with increased risk of developing AD [57, 65], and cause more severe AD-associated neuropathological changes in various mouse models [66, 67]. Here, we observed a significantly increased number of phospho-Tau (AT8)-positive dystrophic neurites around β-amyloid plaques in both lateral entorhinal cortex and hippocampus in mice with A+ and A+T+ genotypes upon TWD. Some studies have reported that HFD or TWD does not have an effect on dystrophic neurites nor Tau pathology in APPswe/PS1dE9 or Tau P301L mice, respectively [20, 22]. However, in an opposite experimental setup, Buccarello et al. showed fewer AT8-positive cells in the cortex and hippocampus of Tau P301L mice fed with low-fat and low-protein diet as compared to STD [9], suggesting that diet can modulate the dystrophic neurite pathology. In the present study, TWD or T+ genotype did not affect β-amyloid pathology in the lateral entorhinal cortex or hippocampus of mice with A+ genotype. This was not surprising, since we have shown previously that β-amyloid burden does not correlate with memory impairment in APPswe/PS1dE9 mice [62]. Furthermore, several studies have reported adverse effects of HFD on memory and learning without exacerbating β-amyloid pathology in AD mouse models [47, 73, 83]. Conversely, other studies have suggested that HFD increases cortical β-amyloid load in APPswe/PS1dE9 [15] and 3xTgAD mice [36]. To our knowledge, our study is the first that demonstrates an increased number of phospho-Tau (AT8)-positive dystrophic neurites around β-amyloid plaques in AD mouse models upon TWD. This is an important finding since dystrophic neurites are a hallmark of dense-core neuritic plaques, which are used in the neuropathological diagnosis of AD owing to their association with cognitive impairment [85].

Studies investigating diet-associated molecular mechanisms in APP23 transgenic mice have shown that HFD increases the expression of genes associated with immune response and inflammation, such as *Trem2*, *Tyrobp*, *P2ry12*, *Dock2* as well as a number of cytokines, chemokines and toll-like receptors [67]. Here, we observed a significant increase in the expression of *Trem2* and *Tyrobp* and other well-established DAM signature genes [40], likewise suggesting an increased immune response in A+Tw and A+T+ mice upon STD. However, it should also be emphasized that A+Tw and A+T+ mice upon STD did not differ in relation to DAM response. Further analysis of the transcriptomic alterations revealed a partially suppressive effect of TWD related to immune response in mice with A+ or A+T+ genotypes. Detailed examination indicated that the genes affected by TWD correlated with *Trem2* expression and were also intimately associated with the DAM signature [40]. This suggests that TWD suppresses the microglial response to β-amyloid pathology and that this repressive response is linked to Trem2-related processes, such as DAM activation. This molecular reprogramming of microglia upon TWD was further supported by the decreased Iba1 and Cd68 positivity around β-amyloid plaques in TWD mice. Thus, both expressional and IHC analyses independently support the idea of a repressed immune response to TWD, which is eventually manifested as decreased number of microglia and increased number of dystrophic neurites around the β-amyloid plaques. Moreover, the decreased intensity of the lysosomal Cd68 may suggest that the functional capacity of the remaining microglia around the β-amyloid plaques is compromised.

Evaluation of the effects of TWD irrespective of the genotype also revealed a substantial overlap with altered expression of the previously described DAM genes. *Pik3r1*, encoding for p85α, was one of the top hits in the DE analysis. Since *Pik3r1* was downregulated in the hippocampus by TWD irrespective of the genotype, we asked whether any key PI3K-Akt signaling pathway components were affected by TWD. Indeed, a decreased phosphorylation of Akt1/2 at the S473/S474 and T305/308/308 activation sites as well as at the inhibitory S9 site of Gsk3β were observed in the hippocampus of TWD mice. Our findings are in line with previous studies showing that the PI3K-Akt-Gsk3β signaling pathway is impaired in mouse and rat brain upon HFD and TWD [33, 73]. Interestingly, previous analyses in human post-mortem samples have demonstrated decreased levels of p85α in the brain of AD patients [63] as well as in AD patients with co-morbid T2D [55]. Importantly, Liu et al. showed that the decrease in the levels of phosphorylated and total p85α was more prominent in the brain of these comorbid patients as compared to AD patients without T2D, and that the alterations in p85α coincided with altered phosphorylation of the downstream targets AKT and GSK3β [55]. Also, previous studies in different mouse models with a diabetic or obese phenotype have detected decreased levels of p85α in the brain tissue [27]. Conversely, caloric restriction in Tg2576 mice led to increased protein levels of p85α in the cerebral cortex as compared to control mice alongside with increased activity of Akt1 [77]. In general, GSK3β is one of the main kinases phosphorylating Tau in the brain [30]. Thus, decreased phosphorylation of the inhibitory S9 site in Gsk3β upon TWD may explain the increased number of AT8-positive dystrophic neurites observed in our study. Supporting this idea, increased phosphorylation of Tau due to HFD or TWD in mice and rats has been observed also previously [4, 33, 73]. On the other hand, our biochemical analysis did not reveal clear changes in the levels of phosphorylated or total Tau by the TWD, which might be due to limited solubility of aggregated Tau protein with the extraction method used. In fact, there are also other studies, in which the effects of HFD or TWD on the phosphorylation status of Tau has not been conclusive in the biochemical assessments [36, 47, 81, 83].

Alterations of PI3K-Akt signaling are known to affect multiple cellular functions, such as autophagy, metabolism, proliferation, cell survival, and growth [44]. Here, we found increased levels of the autophagy markers p62 and LC3BI in the hippocampal total protein lysates of TWD mice, suggesting decreased autophagosomal activity. Given the decreased activation of Akt-kinase due to TWD, this was an unexpected finding since the activated Akt-kinase is known to activate mTORC1, which again inhibits the induction of autophagy [102]. However, our observation related to the increased p62 levels is consistent with previous studies reporting increased p62 levels in the peripheral tissues, hypothalamus, and hippocampus of mice on HFD [13, 75, 104]. Furthermore, HFD has been linked to impaired lysosomal activity [104]. These findings suggest that also other pathways in addition to PI3K-Akt signaling pathway control autophagy. Indeed, 5’ AMP-activated protein kinase (AMPK) is a conserved sensor of nutritional status, which may affect autophagy independently of mTORC1 [42]. Since starvation is a well-known inducer of autophagy, the opposite condition with high energy supply, such as TWD, may result in reduced autophagy.

The fact that *Pik3r1* is abundantly expressed in mouse microglia [107] together with our RNA-sequencing data prompted us to elucidate mouse microglial functions upon downregulation of PI3K-Akt signaling by both pharmacological and genetic approaches. Inhibition of PI3K-Akt signaling using LY294002 both in mouse BV2 and primary microglia significantly decreased the phagocytic uptake of bioparticles in a dose-dependent manner. A similar decrease was observed in the primary microglia isolated from Akt2 KO mice. Interestingly, the treatment of Akt2 KO microglia with LY294002 further reduced the phagocytic uptake of bioparticles, suggesting that PI3K-Akt pathway as a whole entity plays a key role in the regulation of phagocytic activity in microglia. LPS treatment of Akt2 KO microglia cultures reduced the levels of the inflammatory mediators Tnf-α, Il6 and NO in the culture medium, which agrees with the previous findings indicating that Akt2 KO macrophages are hyporesponsive to LPS [2]. Apart from being the regulatory subunit of PI3K, p85α encompasses a significant role in controlling the actin organization and cell migration, independently of PI3K activity [35]. Therefore, it is possible that the reduction in the expression of *Pik3r1* owing to TWD does not only affect PI3K activity, but also other functions that are relevant for microglia. On the other hand, studies in *Pik3r1* knock-out mice have revealed that the lack of p85α subunit reduced the activity of PI3K and Akt in the peritoneal exudate cells upon LPS-induced stress [59]. Moreover, the LPS treatment of *Pik3r1* knock-out mouse cells induced the expression of *Tnf* and *Il6* as well as increased the nuclear localization of phosphorylated transcription factor ATF-2, known to increase proinflammatory cytokine expression in LPS-stimulated monocytes [59]. In conclusion, modulation of the activity of PI3K-Akt signaling pathway affects central functions of microglia, such as phagocytic uptake, autophagosomal degradation, and proinflammatory response against different stressors, including β-amyloid. In this context, TWD-associated reduction in the expression of *Pik3r1* encoding p85α may significantly contribute to the observed suppressive immune response and reduced clustering of microglia around the β-amyloid plaques. On the other hand, the results from the IHC analysis of p85α were partially conflicting, since the area and intensity of p85α around β-amyloid plaques was increased upon TWD in A+Tw mice, while in A+T+ mice, the effect was the opposite. However, this type of genotype dependency is also observed in the RNA-sequencing data, where the most aberrant effects on transcriptional response were observed in A+T+ mice.

Several studies have reported that HFD affects particularly microglial phenotype, function and/or inflammatory outcomes [3, 20, 98]. Trem2 has been shown to play a central role in AD and one of its functions is to act as a lipid-sensing receptor affecting microglial response [100]. In addition to microglia and cells of myeloid origin (e.g. monocytes, macrophages and osteoclasts), Trem2 is also expressed in mature adipocytes [70], and has been shown to affect insulin resistance in HFD mice through adipose tissue remodeling [54]. These reports emphasize that alterations in Trem2 signaling may also mediate the effects of HFD or TWD. Related to this, we now show that the ligand-induced activation of Trem2/Dap12 both directly (Trem2 antibody) and indirectly (M-CSF) upon the inhibition of PI3K-Akt signaling enhanced the activation of Syk kinase in mouse microglia. This was an unexpected result since it has been previously shown that the inhibition of PI3K in macrophages and osteoclasts prevented the recruitment of p85α and Syk kinase to Dap12, leading to reduced downstream signaling via other mediators, such as Plcγ2 [72]. Whether the observed increase in Syk kinase activation is a compensatory effect for the reduced downstream signaling in microglia remains to be determined in future studies. Collectively, these results highlight the seminal role of p85α/PI3K in the activation of Trem2/Dap12 signaling pathway, particularly in the formation and activation of Syk kinase complex in microglia. Recent studies suggest that impaired Trem2-Akt-mTOR signaling in AD patients carrying *TREM2* risk variants and in Trem2-deficient mice with AD-like microglia affects autophagy and metabolism [95] and decreases the ability of microglia to form a protective barrier around β-amyloid plaques, leading to axonal dystrophy [106]. The findings of the present study, including the increased number of dystrophic neurites and impaired PI3K-Akt signaling, large range of transcriptional changes in the microglial genes due to TWD and/or A+ genotype, and decreased Iba1 and Cd68 immunoreactivity around β-amyloid plaques similarly to previous studies, suggest that indeed impaired PI3K-Akt signaling reduces microglia clustering and their ability to form a protective barrier around plaques [95, 106]. Consequently, this may lead to leakage of toxic β-amyloid aggregates disturbing neuronal integrity. Our observation that TWD increased the number of AT8-positive dystrophic neurites around the β-amyloid plaques supports this hypothesis. RNA-sequencing did not reveal major alterations in the expression of *Gfap*, suggesting that TWD does not affect astrocyte reactivity.

In order to elucidate whether similar alterations to those observed in microglia in the TWD mice could also be detected in human brain tissue, we utilized frontal cortical brain biopsies obtained from iNPH patients harboring β-amyloid pathology. Brain biopsies taken during shunt surgery offer a unique window into the brain tissue of living iNPH patients to examine the early stages of AD-related brain pathology [53]. We found that the combination of obesity and T2D led to presence of fewer microglia around the β-amyloid plaques compared to normal weight and obese iNPH patients without T2D, which is well in concordance with our findings in mice with the TWD-induced diabetic phenotype. Neuropathological examination revealed no Tau pathology (AT8 immunoreactivity) in the cortical tissue samples from the iNPH patients with respect to normal weight, obese, or diabetic phenotype, suggesting that β-amyloid drives the microglial alterations in the brain of these patients. Altered microglial response to β-amyloid plaques due to T2D has been suggested earlier, although no quantitative data have been presented [96]. Interestingly, increased microglial dystrophy has been reported in the hypothalamus of obese individuals [3] as well as in the hippocampus and several cortical areas in AD patients [76, 91]. Collectively, our results together with previous studies in both mouse and human support the idea that obesity and T2D contribute to microglial dysfunction. According to our present data, particularly T2D-related metabolic changes appear disturb microglial function around β-amyloid plaques. We have focused here on the most prominent effects of TWD in mice with different AD- and/or tauopathy-associated genetic backgrounds. However, the identified novel targets and regulatory networks associated with the individual genotypes (AwT+, A+Tw, and A+T+) will set the basis for further elucidation of the underlying mechanisms related to the comorbidity of AD and T2D. Moreover, the fact that similar results were obtained in the brain tissue of TWD mice and iNPH patients with T2D lays an outstanding foundation to test novel therapeutic interventions, focusing specifically on the microglial response and clustering, against AD-associated pathologies in the brain.

## Acknowledgements

This study was supported by Academy of Finland (grant numbers 288659, 307866, 315459); Sigrid Jusélius Foundation; the Strategic Neuroscience Funding of the University of Eastern Finland; JPco-fuND2 Personalised Medicine for Neurodegenerative Diseases (grant number 334802); Doctoral Programme in Molecular Medicine (DPMM) at the University of Eastern Finland; European Union Horizon 2020 research and innovation programme (grant number 692340);Finnish Functional Genomics Centre, University of Turku and Åbo Akademi; and Biocenter Finland. Part of the work was carried out with the support of UEF Cell and Tissue Imaging Unit, University of Eastern Finland, Finland. The computational analyses were performed on servers provided by UEF Bioinformatics Center, University of Eastern Finland, Finland.

