## Supplemental material and figures for "Diabetic phenotype in mouse and humans with β-amyloid pathology reduces the number of microglia around β-amyloid plaques"

**Supplementary Figure Legends**

**Supplementary Fig. 1 a-c** Assessment of soluble Aβ42 and Aβ40 levels as well as Aβ42 and Aβ40 ratio of hippocampal samples from mice with A+ genotype revealed no significant diet effect. Data are presented as mean + SEM, n=5-6, Two-way ANOVA. **d** A representative Western blot image of hippocampal lysates and quantification phospho-Tau and total Tau levels. Phosphorylated protein levels were normalized to their respective total protein levels in cell lysates and total protein levels were normalized to β-actin. All results are shown as mean + SEM, n=5-7 mice/group, Two-way ANOVA.

**Supplementary Fig. 2 a** Heatmap of z-score values for transcripts specifically mapping to human (h) *APP*, *PSEN1*, and *MAPT,* show a significant increase in *APP* and *PSEN1* in A+Tw and A+T+ mice, and a significant increase in *MAPT* levels for AwT+ and A+T+ mice as compared to AwTw mice. **b** PCA for all samples (AwTw, AwT+, A+Tw, A+T+ STD/TWD). **c** PCA for samples of each genotype showing segregation due to TWD. **d** Number of DE genes (FDR < 0.05) for each genotype/diet combination as compared to corresponding diet AwTw mice. **e** Venn diagram showing overlap of DE genes between genotype/diet groups. n=4-6 mice/group.

**Supplementary Fig. 3 a** Heatmap of z-score values for genes with Z > 2.5 or < -2.5 in figure 2A for STD mice, but not TWD mice for either A+Tw vs AwTw mice (left) or A+T+ vs AwTw mice (right). **b** Correlating and hierarchical clustering of modules in STD or TWD networks. **c** Cell type enrichment of genes in network modules. **d** Box plots representing eigengene values for modules negatively associated with genotype. Points represent eigenegene values for each mouse in that group. Box plots show the median, 25th and 75th percentiles, error bars show 1.5 interquartile ranges. n=4-6 mice/group.

**Supplementary Fig. 4 a** *Pik3r1, Trem2*, and *Tyrobp* expression levels obtained from RNA-sequencing, and **b** from qPCR

**Supplementary Fig. 5 a** Quantification of Western blot shown in Fig 5 revealed that 10 μM LY294002 together with LPS increased LC3B II levels as compared to both untreated and LPS-treated controls (p<0.05, One-way ANOVA, LSD posthoc). **b** LY294002 treatment with or without LPS had no effect on LC3B I levels (Welch's ANOVA, Tamhane's posthoc). **c** Quantification revealed increased total Akt level in samples treated together with LPS and 1 μM LY294002 as compared to untreated samples (UNT, $p<0.05$, Welch's ANOVA, Tamhane's posthoc) **d** Quantification of p85α levels showed significantly decreased levels in samples treated together with LPS and 10 μM LY294002 as compared to both UNT and LPS treated control samples ($p<0.05$, Welch's ANOVA, Tamhane's posthoc). **e** A representative Western blot image from lysates of BV2 cells treated with LY294002 (LY) (0.1 μM – 10 μM), and **f** with LY294002 together with LPS. **g** Quantification revealed decreased phosphorylation status of Erk1 and 2 (pErk1 and 2) in Y204 site in a dose-dependent manner in samples treated with LY294002 or treated together with LPS and LY294002 (One-way ANOVA, LSD posthoc). **h** Quantification of total Erk2 levels showed an increase in cells treated with 0.1 μM and 1 μM LY294002 together with LPS (One-way ANOVA, LSD posthoc). **i-j** Quantification revealed no changes in pro-caspase-3 or in active caspase-3 levels (Welch's ANOVA). Phosphorylated protein levels were normalized to their respective total protein levels in cell lysates and total protein levels were normalized to β-actin. Data are presented as mean + SEM from four biological replicates. *$p<0.05$, **$p<0.01$, ***$p<0.001$ vs. untreated control, #$p<0.05$, ##$p<0.01$, ###$p<0.001$ vs. LPS control.

**Supplementary Table 1.** Differential expression analysis results for AwT+, A+Tw, and A+T+ vs AwTw, SD mice

**Supplementary Table 2.** Differential expression analysis results for AwT+, A+Tw, and A+T+ vs AwTw, TWD mice

**Supplementary Table 3.** Enrichment analysis for differentially expressed genes in AwT+, A+Tw, and A+T+ vs AwTw comparison for both SD and typical Western diet mice

**Supplementary Table 4.** Enrichment analysis for genes showing discordant response to amyloid- and/or tau-pathology upon TWD (figure 2A, blue dot genes)

**Supplementary Table 5.** WGCNA module assignment and membership values

**Supplementary Table 6.** Enrichment analysis for WGCNA module genes

**Supplementary Table 7.** Differential expression analysis results for TWD vs STD mice

**Supplementary Table 8.** Enrichment analysis for differentially expressed genes upon TWD

### Supplementary Materials & Methods

*Real time quantitative PCR analysis*

RNA was isolated from mouse hippocampus homogenates using the Direct-zol RNA MiniPrep (Zymo Research, Irvine, CA, USA). RNA concentrations were measured using the NanoDrop ND-1000 spectrophotometer (Thermo Fisher Scientific, Waltham, MA, USA). Total of 250 ng of RNA was subsequently synthesized into cDNA using the Transcriptor First Strand cDNA Synthesis Kit (Roche Diagnostics, Risch-Rotkreuz, Switzerland). RT-qPCR was subsequently run using the LightCycler 480 Instrument II (Roche Diagnostics, Risch-Rotkreuz, Switzerland) with the LightCycler 480 SYBR Green I Master (Roche Diagnostics, Risch-Rotkreuz, Switzerland) and the following primers (TAG Copenhagen, Copenhagen, Denmark): mouse Pik3r1 forward 5'-TATTGCGAGGGAAGCGAGAC -3', mouse Pik3r1 reverse 5'- ACTTCGCCGTCTACCACTAC-3', mouse Trem2 forward 5'-TGG AAC CGT CAC CAT CAC TC-3', mouse Trem2 reverse 5'-TGG TCA TCT AGA GGG TCC TCC-3', mouse Tyrobp forward 5'-ACC CGG AAA CAA CAC ATT GC-3', mouse Tyrobp reverse 5'-TTG CCT CTG TGT GTT GAG GT-3', mouse β-actin forward F: 5'-GGCTGTATTCCCCTCCATCG-3', mouse β-actin reverse R: 5'-CCAGTTGGTAACAATGCCATGT-3'. Results were calculated using the 2–ΔΔCt method (Livak and Schmittgen, 2001).

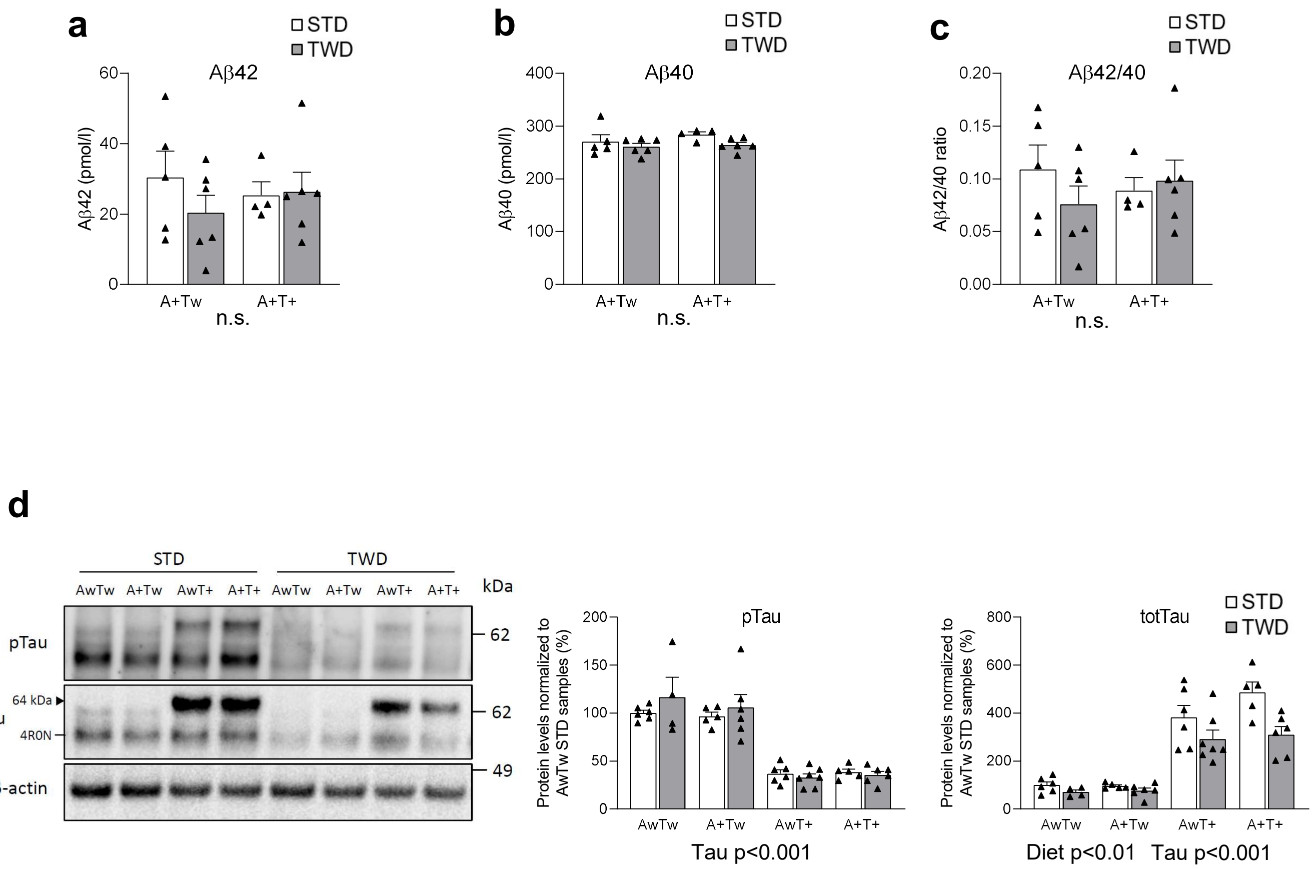


**Supplementary Figure 1**

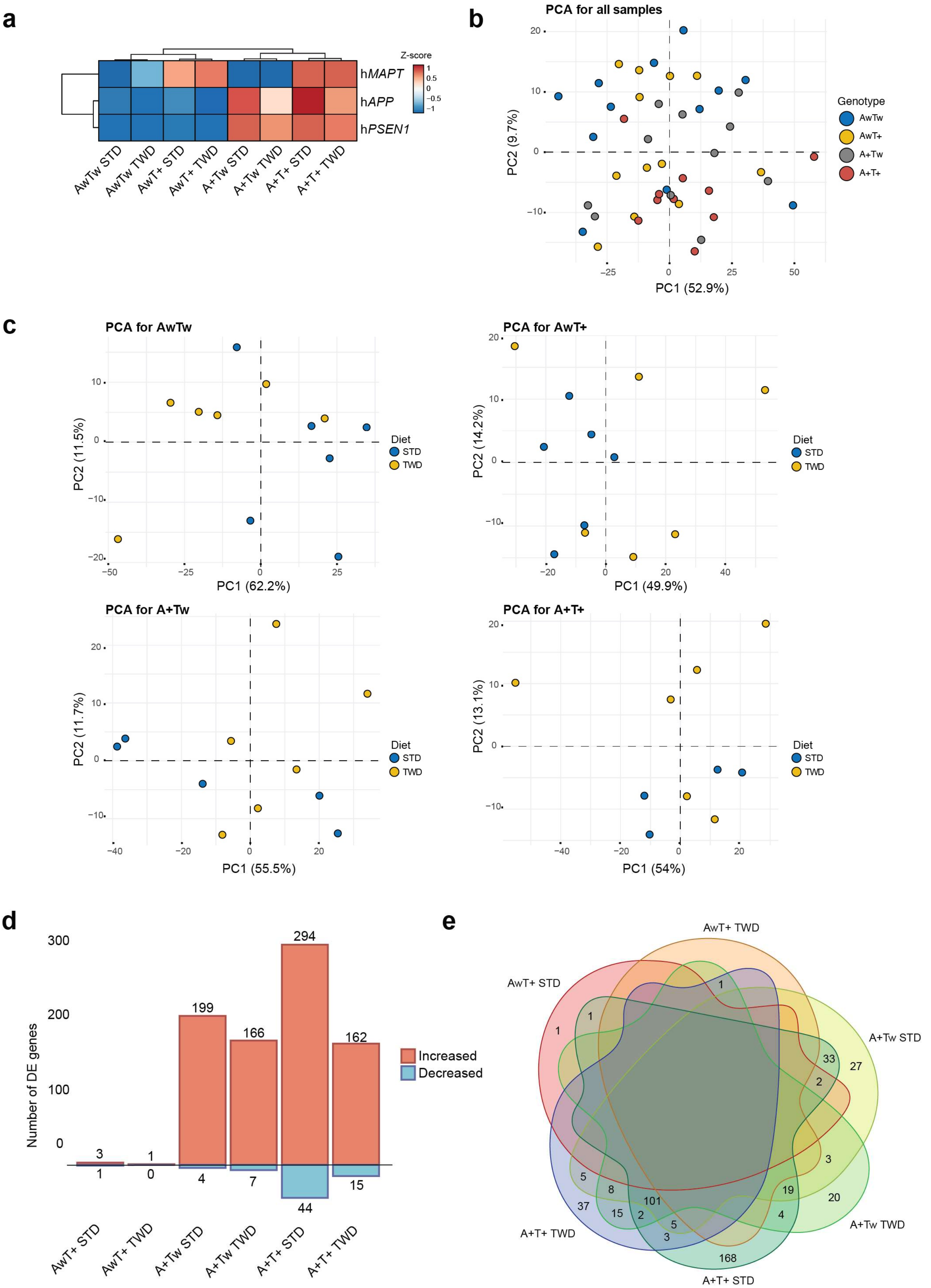


**Supplementary Figure 2**

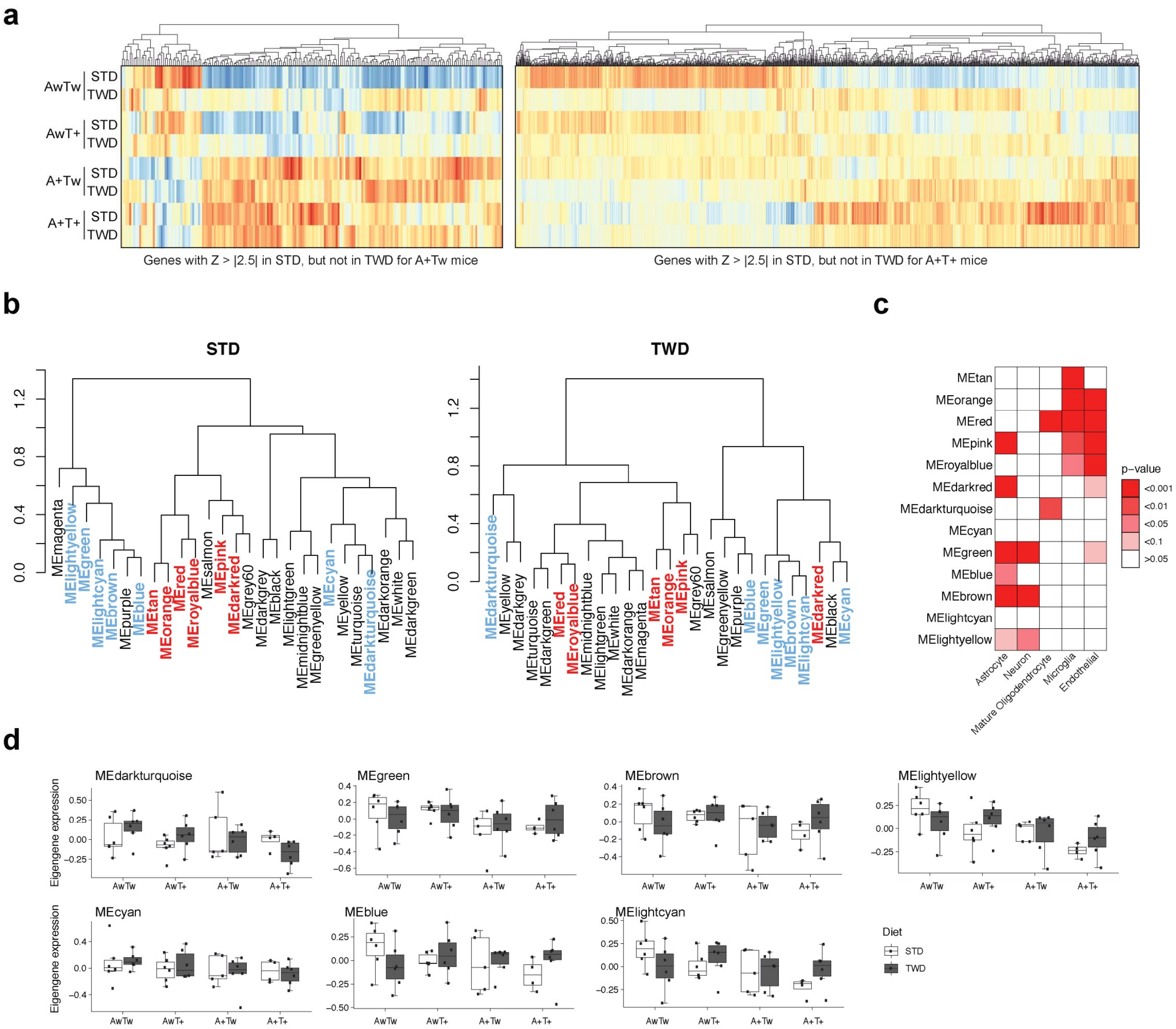


**Supplement Figure 3**

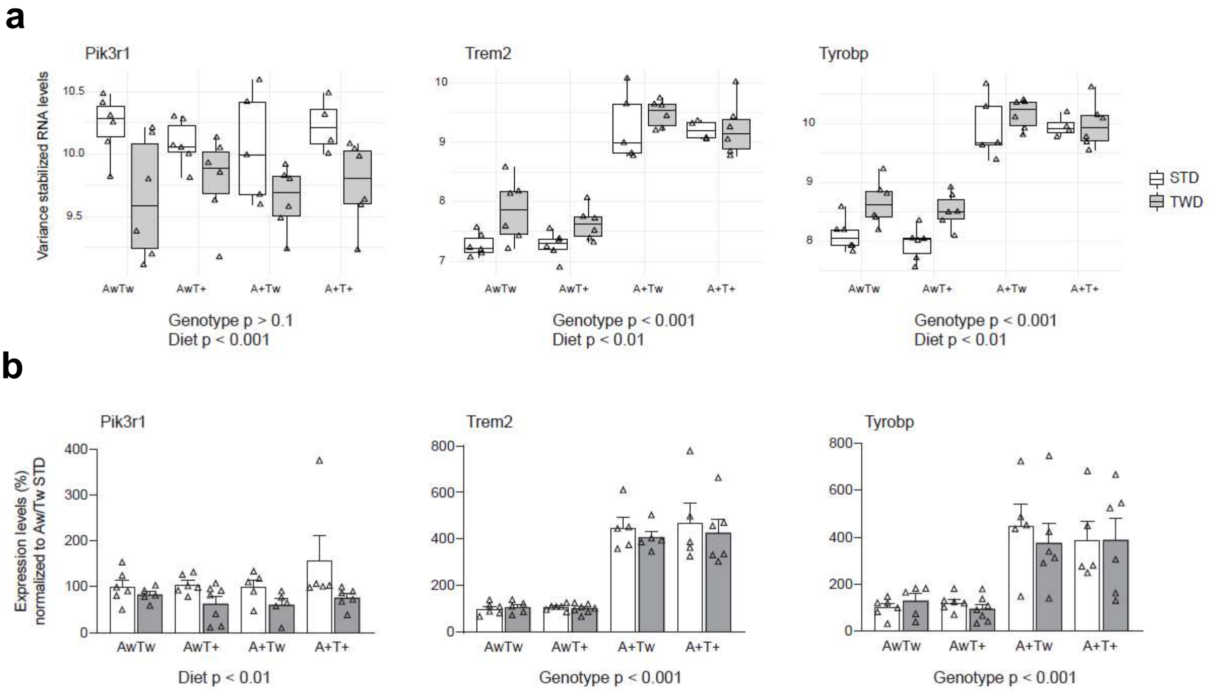


**Supplement Figure 4**

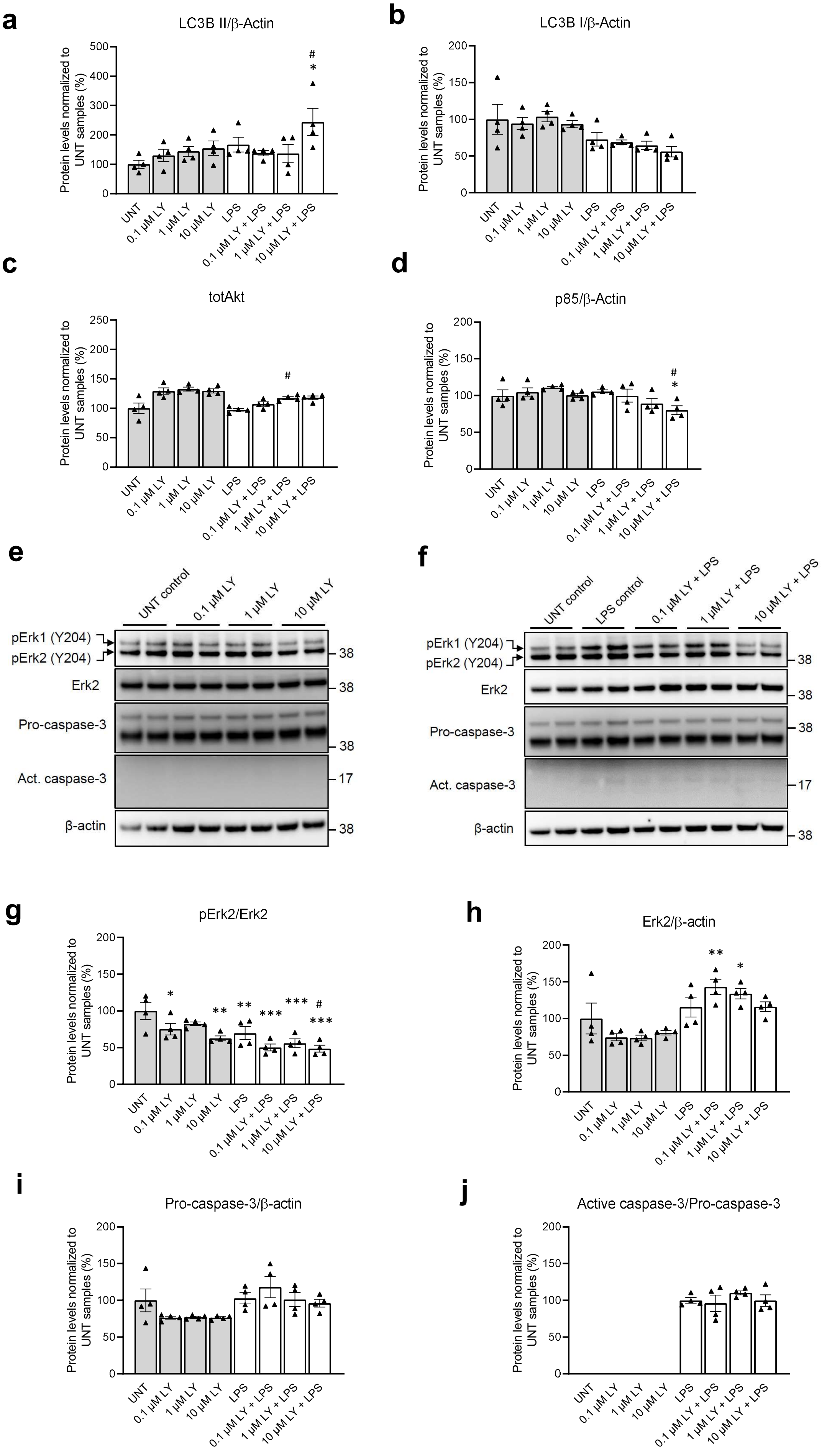


**Supplement Figure 5**
